# Designed IDPs phase separate and mix or demix according to sequence designed parameters

**DOI:** 10.64898/2026.09.23.753891

**Authors:** Arjun Singh, Gregory L Dignon

## Abstract

Biomolecular condensates are condensed assemblies of biomolecules that form thrwough the process of liquid phase separation. Condensates in biology typically function as membraneless organelles, providing compartmentalization in the absence of a dividing lipid membrane. The molecular make-up of different condensates includes diverse multivalent proteins and nucleic acids as the primary drivers, many of them including significant fractions of intrinsically disordered regions (IDRs). To date, the sequence to phase separation relationship of IDRs has focused largely on one protein at a time, studying single-component condensate formation, or single-component partitioning into condensates. The co-phase separation and mixing of two or more IDRs is considerably more complex as both sequences can vary widely in their self- and cross-interactions, as well as their relative abundance in solution. It is unclear that a rule which predicts how a single sequence behaves will also predict what happens when two sequences are mixed. In this study, we disentangle the influence of sequence from that of composition using a set of 18 LAF-1 RGG variants that keep the same length and amino-acid composition and change only the order of the residues. This lets us vary charge patterning and, to a lesser extent, hydropathy patterning while keeping protein composition fixed. By themselves, the sequences phase separation and single-chain compaction are controlled by their degree of charge and hydropathy patterning. Within single-component condensed phases, each sequence adopts a more extended conformational ensemble, due to a more favorable, self-solvated environment. We find that mixing two IDRs together into a condensate causes this universal scaling behavior to break, impacted by the relative interactions of the two components and overall composition of the slab. We find two different qualitative behaviors, one characterized by cooperative co-condensation when both sequences are subcritical, and the other by scaffold–client behavior when one sequence is supercritical. The scaffold-client systems generally show a high degree of demixing, while the co-condensing systems are generally quite well-mixed in the dense phase. This is surprising because even in cases where both partners have significantly different patterning parameters, they still mix. Thus, the descriptor that predicts a sequence’s behavior alone can help indicate whether it will mix with or separate from a second component, but it does not fully determine the outcome on its own.

## Introduction

Cells organize much of their biochemistry through compartmentalization without a membrane. This is accomplished by certain proteins and nucleic acids, which can demix from the surrounding cytoplasm or nucleoplasm into dense, liquid-like droplets, called biomolecular condensates [1, 2]. These condensates are found throughout the cell, including P granules, stress granules, the nucleolus, and heterochromatin, and many others [3–7]. Condensates can selectively concentrate specific molecules, accelerating or slowing particular reactions and enabling cells to support natural growth and replication, as well as responding rapidly to stress[8–12]. When condensate formation is disrupted, it may also contribute to disease, including several neurodegenerative disorders and some cancers [4, 13–17].

One common mechanism of biomolecular-condensate formation is liquid–liquid phase separation (LLPS), in which proteins and nucleic acids condense into a dense, liquid-like phase that coexists with a surrounding dilute phase [3, 18]. Many proteins that drive LLPS contain intrinsically disordered regions (IDRs), which are segments of the protein chain that do not have a single stable structure [19–21]-. Because IDRs remain conformationally flexible, they can present multiple interaction motifs and form many weak, reversible contacts within and between molecules [21–23]. Purified IDRs and multidomain proteins can undergo LLPS through multivalent favorable interactions, whereas many isolated globular proteins lack sufficient valency to condense [18, 22, 24, 25]. An IDR’s phase behavior is encoded by its amino-acid sequence, including residue composition, charge and hydropathy patterning, and the identity and distribution of interaction motifs [26–30]. This provides a path for linking genome-encoded sequence features to condensate behavior in cells [31–33].

Two key sequence features are composition and patterning [34, 35]. Composition is simply which amino acids are present and in what fraction, for example how many residues are charged, aromatic, hydrophobic, etc. or how many of a particular amino acid are present, like special cases glycine and proline[32]. Patterning refers to how those residues are arranged along the sequence of amino acids. A charged residue can sit in the vicinity of other charged residues of the same sign, forming a block, or it can be spread evenly among opposite charges [36]. Metrics like the sequence charge decoration (SCD) and the sequence hydropathy decoration (SHD) were built to capture this kind of patterning for charged residues, and hydrophobic/aromatic clustering [37, 38]. Previous work has shown that sequence patterning can influence single-chain compactness, condensate density, and saturation concentration, even among variants with identical amino-acid compositions [29, 39–42].

While relatively simple patterning properties do a decent job of capturing single-chain properties, and single-component LLPS behavior, cellular condensates are rarely composed of a single protein[43, 44]. P bodies and stress granules are multicomponent ribonucleoprotein assemblies that can contain hundreds of proteins along with diverse RNA molecules [45–48]. To study multicomponent condensate mixing under controlled conditions, researchers reconstitute condensates in vitro from a small number of purified IDR-containing proteins and, when relevant, RNA [5, 49]. A reasonable conclusion is that whether two types of molecules occupy the same droplet, form distinct and separate droplets, or organize into distinct nested, or attached droplets depends largely on their interactions with one another, in addition to each one’s self-interactions[50–52].

On the computational side, emerging tools are making multicomponent IDR questions more tractable. FINCHES predicts the net chemical specificity between two IDR sequences directly from sequence, without requiring molecular simulations [53, 54]. A related sequence-based approach has used the same kind of reasoning to predict which combinations of components will stay mixed and which will demix into separate condensates [55]. Once a two- or multicomponent condensate has formed, a separate question is how to quantify the extent to which its components are mixed. A statistical potential was also developed by Villegas and Levy which reports a stickiness scale with promise to predict partitioning of general proteins into condensates[56]. A recently developed spatial order parameter, the demixing index *D*_demix_, divides the condensate into small voxels and compares each voxel’s local composition with the overall composition [57]. *D*_demix_ is near zero for uniform mixing and increases as the components become spatially segregated. Together, these methods give us ways to predict pairwise interactions and to quantify the resulting spatial organization once a condensate has formed.

What these tools do not yet tell us is whether the same simple sequence descriptors that govern single-component phase behavior also predict mixing in two-component systems. This is hard to test with natural sequences, since composition and patterning almost always vary together. Towards this, we design a controlled test, using sequences of identical length and composition that differ only in residue arrangement, studied first alone and then in mixtures. We use a previously designed library of 18 LAF-1 RGG-domain variants with identical lengths and amino-acid compositions but different charge arrangements [30]. We confirm that charge patterning alone, measured by SCD, controls how compact each variant is and how readily it phase separates on its own. We then mix each variant with a fixed reference sequence, LAF-1 V2, and use the demixing index to test whether the two proteins mix, partially separate, or largely exclude each other. Mixing appears to be only minimally influenced by the difference in SCD between partners. Instead, variants that phase-separated readily on their own generally mixed with V2, whereas weakly self-associating variants were largely excluded. We finally demonstrate how single-sequence descriptors are a useful guide to predict single-component phase behavior, but not enough on their own to predict mixing or demixing. This composition-matched framework can be extended to other sequences and experimental systems, providing a potential design rule for engineering condensates with defined composition and organization [54].

## Results and discussion

### Sequence order affects chain size and phase behavior

Intrinsically disordered regions (IDRs) are known to be significant drivers of biomolecular condensates formation [3, 25]. Their behavior depends not only on which amino acids they contain but also on how those amino acids are arranged along the chain [27, 29, 36]. Separating these two effects is difficult, because most natural sequences differ in both composition and order at the same time. To understand the effect of amino-acid arrangement, we used a library of 18 sequences derived from the LAF-1 RGG region that we built in our previous study [30]. Every version had 168 residues and exactly the same amino-acid composition, the only difference being the order in which those residues appeared along the sequence. To describe the sequence patterning, we use sequence charge decoration (SCD) and sequence hydropathy decoration (SHD). SCD decreases as similarly charged residues become more blocky, whereas SHD reflects the degree of hydrophobic-residue clustering [37, 38]. The library scans SCD from about 0 (well-mixed, close to the wild type) to about −21 (strongly blocky) while keeping SHD within a narrow band (Figure 1A). The library is limited to mostly negative SCD values since LAF-1 RGG has nearly zero net charge.Essentially all designed variants have residue reordering which made the charges more segregated or blocky. This gives a controlled set in which charge and hydropathy patterning are the main controlling variables, so we can ask how patterning alone shapes the behavior of a single chain and of the condensed phase it forms. Previously we found that for these sequences, SCD had a majority influence on phase separation behavior, while SHD had only a moderate influence, though this would likely not be the case for sequences that have fewer charged amino acids[30, 38].

**Figure 1:**
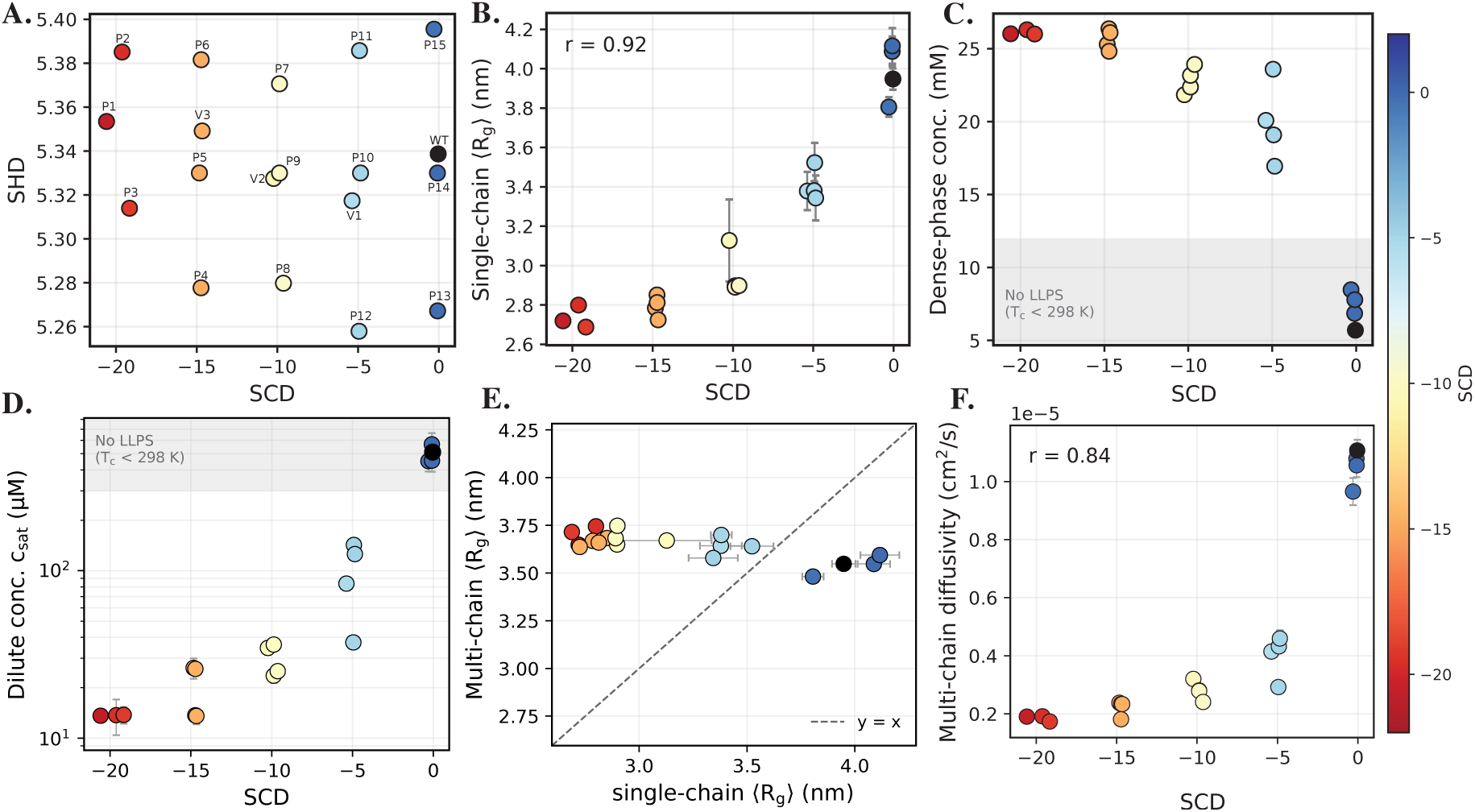
Charge patterning controls single-chain dimensions and single-component phase behavior in a composition-matched LAF-1 sequence library. A) Design space of the LAF-1 RGG wild type and 18 derived sequences (19 systems in total) in the plane of SCD and SHD. B) Mean single-chain radius of gyration *<R_g_>* from single-chain simulations at 298 K versus SCD. C) Dense-phase concentration of the single-component slab simulations versus SCD. D) Dilute-phase saturation concentration *c_sat_* versus SCD on a logarithmic axis. E) Single-chain *<R_g_>* versus Multi-chain *<R_g_>* from the slab simulations. F) Chain translational diffusivity D in the slab versus SCD.

As a first step, we analyzed each sequence individually. At a fixed amino-acid composition, charge patterning has a clear, systematic effect on chain size. The average radius of gyration declines from about 4.1 nm in well-mixed sequences to about 2.7 nm in highly blocky sequences, exhibiting a strong relationship with SCD (Figure 1B). In contrast, hydropathy patterning shows almost no relationship with chain size across the same library (see SI Figure 2A). Together, these results indicate that, at fixed composition, singlechain compactness is influenced more by charge patterning than by hydrophobic-residue patterning in this sequence library.

We then ran multi-chain slab simulations to measure the phase behavior of each sequence on its own, using the HPS-Urry model and slab simulation methodology as in previous work [30, 58]. The concentration of the dense phase increases as the sequence becomes more charge-segregated, from about 6 mM for the well-mixed sequences to about 26 mM for the blockiest (Figure 1C). We note that the low values of concentration 12*mM* indicate a sequence that does not form a stable slab in the simulation, and thus we are calculating the local density of any large clusters in the simulation box for those cases. We calculated the dense- and dilute-phase concentrations using the domain-decomposition method (DDM) developed previously by Morton et al. [57]. We divided the simulation box into cubic voxels with edges of about 35 Å. We then used the residue-density distribution across the voxels to identify the dense phase inside the slab and the dilute phase outside the slab. We used a fixed temperature of 298 K for all sequences so that they could be compared under the same conditions. At 298 K, blocky sequences are further below their known *T_c_* values (SI Figure 1) and form denser condensates with lower *c_sat_* values (Figure 1D). The four well-mixed sequences near *SCD* ≈ 0 (WT, P13, P14 and P15) did not phase separate since they were supercritical at 298 K (SI Figure 1). Although these sequences do not show LLPS alone at 298 K, we included them to test whether V2 can promote phase separation. These sequences are shown in the shaded “No LLPS” region in Figure 1C and D. The dilute-phase concentration coexisting with the dense phase, decreases by more than an order of magnitude with increasingly negative SCD (Figure 1D). Thus, we find that generally sequences with stronger charge segregation form more compact chains, have denser condensates and lower saturation concentrations consistent with previous work [29, 36, 39].

Comparing the two ensembles shows that single-chain size in isolation can differ from chain size within the condensate. Both values are single-chain radii of gyration, *R_g_*, which describe the size of an individual chain, but in different contexts. Comparing them shows how the crowded condensate environment changes chain size. The isolated chains span a range of about 2.7 to 4.1 nm, in the multi-chain simulations every sequence adopts a similar size of about 3.5 nm (Figure 1E. Similar trends were reported in earlier studies [36, 59]. This means that the compaction seen for blocky sequences as isolated chains does not carry over to the crowded environment of the condensate, where intermolecular contacts appear to expand the chains to a similar extent regardless of sequence. We also note that the internal scaling exponent *ν* of the chains in the slab is about 0.53-0.55, across the whole library, very close to ideal scaling behavior (SI Figure 2B). This reinforces the observation that chains behave consistent with near-ideal-chain behavior in a condensate, as the surrounding environment of an IDR is very similar to itself[60, 61]. Sequences with very different sizes as isolated chains become more similar inside a single-component condensate. We note that for sequences which do not phase separate, there appears to be a similar Rg-buffering effect which causes collapse of the chains, likely due to intermolecular interactions making the environment closer to an ideal solvent, rather than a good solvent (Figure 1E). This suggests that the dense environment reduces the differences between their conformations. The next sections test whether this is also true when different sequences are present in the same condensate.

**Figure 2:**
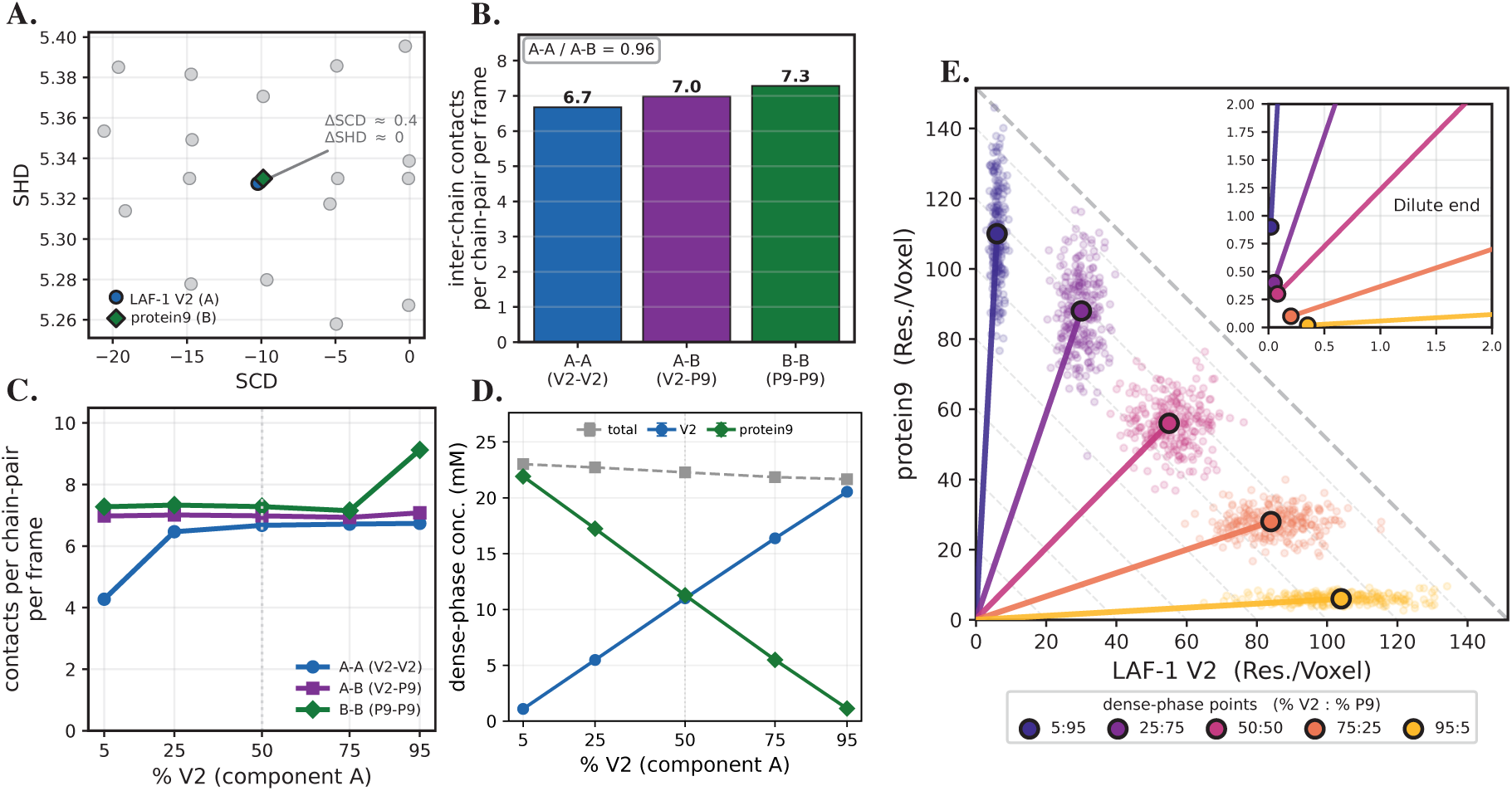
Two sequences with near-identical patterning form a single homogeneous condensate. A) The mixed pair on the SCD–SHD map highlighting V2 and P9 B) Mean intermolecular contacts per chain-pair per frame in the equimolar mixture, resolved into the three block types. C) Contact strength versus composition. D) Dense-phase concentration versus composition. E) Phase diagram shown using the domain-decomposition, per-voxel method. All error bars show block-averaged SEM.

Finally, charge patterning also tracks the dynamics of the condensed phase. The translational diffusivity of chains decreases with charge segregation, from about 1.1 × 10*^−^*^5^ cm^2^*/*s for the well-mixed sequences to about 1.8 × 10*^−^*^6^ cm^2^*/*s for the blockiest (Figure 1F), and the radius of gyration relaxation time (*τ_Rg_*) rises over the same range, from about 1 ns to about 9 ns (SI Figure 2C). The denser condensates formed by blocky sequences are therefore also the slower, more viscous ones. This connection is direct: diffusivity decreases almost monotonically with dense-phase concentration across the whole library (SI Figure 2D). The complete set of single-chain and single-component, multi-chain properties for all 19 sequences is listed in SI Table 1, along with block-averaged standard errors.

Taken together, Figure 1 shows that for our library of maximally modified sequences of identical composition, charge patterning is a strong and consistent predictor of single-chain size, dense-phase concentration and saturation concentration, while internal dynamics are strongly influenced by the dense-phase concentration. Single-chain scaling inside the dense phase seems largely independent of density, or patterning of amino acids. This is a useful starting point, but it describes each sequence only on its own. Biomolecular condensates in cells are generally mixtures of many different molecules, and the way multiple different sequences interact when mixed is not trivially related to how each behaves separately.

### Near-ideal mixing with a similar sequence

Two sequences that phase-separate similarly on their own can still mix uniformly, partly demix, or exclude one another, depending on the balance between the interactions they make with themselves and the interactions they make with each other [5, 50, 52]. Because our library holds composition fixed and varies only patterning, it is well suited to test this directly: the sequences share the same building blocks but differ in self-interaction strength, so mixtures of them isolate the effect of that mismatch. We thus selected a scaffold protein toward the center of our sequence patterning space (i.e. LAF-1 V2; Fig. 1), and tested its mixing behavior with different stoichiometric ratios of each of the other designed sequences, as discussed in the following sections. We first consider the case where two sequences have very similar patterning properties, using LAF-1 V2 with P9, a composition-matched variant whose charge and hydropathy patterning are essentially identical to V2’s (Δ*SCD* ≈ 0.4, Δ*SHD* ≈ 0; Figure 2A). In the SCD–SHD map of Figure 1, P9 and V2 nearly overlap. Importantly, both of these sequences are sub-critical at 298 K and would phase separate individually with a similar propensity. This makes P9 a useful test case for whether similarity in sequence patterning predicts co-assembly with V2. We simulated five different mixtures of these two components at varying stoichiometric ratios for the total composition at 298K.

We first examined the individual intermolecular contact types in the 50:50 mixture, which directly supports the hypothesis of ideal mixing behavior having nearly identical number of like and unlike interactions between chain pairs (Fig. 2B). As shown in the residue–residue contact maps in SI Figure 4A-C, all three contact types mainly involve the charged N- and C-terminal blocks. Importantly, these are normalized by the total number of possible chain-chain pairs of each type, which enables us to repeat this analysis for other stoichiometries besides the equimolar case. When looking across compositions, we find that the three contact densities are equal to within a few percent except at the highly nonstoichiometric cases (Fig. 2C). This contact pattern reflects near-ideal mixing between the two components. At each well-sampled composition (25%, 50%*, and* 75%) the contact counts stay near 7, indicating that the same mixing pattern persists throughout this range (Figure 2C).

At the composition endpoints, the V2–V2 contact count depends strongly on the number of available V2 chains. Self interactions for V2 fall to 4.3 when it is in the minority and the P9 self interactions rise to 9 when it is in the minority. Thus, minority V2 chains make fewer self-contacts, while minority P9 chains make more. Part of this is simply statistics, since with only five minority chains there are just ten same-type pairs to average over, so these endpoint values are noisier than the rest. However, P9–P9 contacts are consistently slightly higher than V2–V2 contacts at all compositions indicating a slightly higher self-interaction propensity/stickiness. At low P9 concentration, the few P9 chains appear to cluster instead of mixing evenly throughout the V2-rich condensate SI Figure 4. However, this observation is based on only five P9 chains in a single trajectory. It is therefore a weak signal. Whether the chains remain together, separate from one another, or localize near the interface may depend as much on their initial placement and stochastic motion as on a real tendency to self-associate.

When looking at the total condensate composition via the dense-phase concentrations, we also observe nearly ideal mixing, having each component closely follow a linear trend along with its total system mole fraction (Figure 2D). Thus, both proteins are recruited into the dense phase in proportions that match their overall system composition. Neither protein is concentrated or excluded relative to its starting amount, so the condensate behaves as one mixed phase that follows roughly the same relative composition between the two components as the overall system with slight variations due to the slightly stronger interactions of P9 compared to V2. The total protein concentration in the dense phase, including both V2 and P9, stays nearly constant across all mixture ratios, reflecting a linear combination of the single-component condensate densities of each component. This behavior is similar to ideal mixing which is what we expect for these two proteins with similar patterning properties and single-component phase behavior.

Figure 2E shows the phase behavior using the DDM voxel method with 35 Å voxels. The scatter plot points show the V2 and P9 densities in individual voxels. The tie-lines connect the dilute phase near the origin to the dense phase, shown by filled circles. The filled circles represent the average composition of the condensate core. The dense phase has nearly the same V2-to-P9 ratio as the starting mixture. At 25:75, 50:50, and 75:25 mixtures, the dense-phase ratios are about 0.34, 0.98, and 3.0, reflecting a near-ideal mixing and co-phase separation with neither component preferentially incorporated. Again, we find that the total dense-phase density remains similar across different compositions, at about 110–118 residues per voxel.

When looking at local densities, we find that both proteins are present in the same dense slab at every composition SI Figure 4A. Even when one protein makes up only about 5% of the system, it still forms a small density peak in the center of the slab. This shows that the minority protein remains in the condensate instead of being excluded. Its dense-phase concentration is about 6 residues per voxel at the most uneven mixtures. Outside the slab, both proteins have densities below 1 residue per voxel, and forming a convex low-density arm. We further quantify the degree of mixing using the demixing index which measures whether the two proteins separate spatially rather than simply change their contact balance using the spread of scatter points with respect to the major axis, roughly following the tie lines[57]. A value of *D*_demix_ = 0 indicates that the two proteins are mixed throughout the condensate, whereas *D*_demix_ = 1 indicates that they occupy separate regions [57]. In this case, its value stays close to zero (*D*_demix_ ≤ 0.033), which is well below the previously-defined mixed-state cutoff of *D*_demix_ *<* 0.2 SI Figure 4B.

The two proteins also match in their basic physical properties across the mixtures (SI Figure 3D–F). Their single-chain sizes are nearly the same, with *R_g_* values of about 37 Å for both proteins, and similar to the values found for single-component condensates, remaining relatively unchanged with different mixture compositions (SI Figure 3D). The internal scaling exponent is also nearly identical for both proteins, with *ν* ≈ 0.552 (SI Figure 3E). This shows that both remain similarly expanded inside the condensate. They also diffuse at similar rates, around 3×10*^−^*^6^ cm^2^ s*^−^*^1^ (SI Figure 3F). Overall, chain size, chain shape, and diffusion do not distinguish V2 from P9 in a mixed condensate. The main difference appears only in self-contacts at strongly uneven mixtures. P9 shows slightly more P9–P9 contacts than V2 shows V2–V2 contacts, especially when P9 is the minority. This suggests a weak preference for P9 self-association. Since *R_g_* and *ν* remain unchanged, this effect likely reflects interactions between different chains rather than chain collapse.

### Mixing with a highly self-attractive protein

We then ask how phase separation and mixing of V2 is influenced by a partner that differs significantly in its patterning.P2 is one of the most charge-segregated sequences in the library, which makes it a strong self-associator, having a critical temperature of 328.67 K (SI Figure 1). Its SHD is also slightly higher than that of V2. On the SCD–SHD map it is located well away from V2, about nine SCD units off, and at a higher SHD (Figure 3A). This partner differs significantly from V2 in charge patterning, shifting toward greater charge segregation, so it is an optimal test of whether sequence difference on its own pushes two proteins apart.

**Figure 3:**
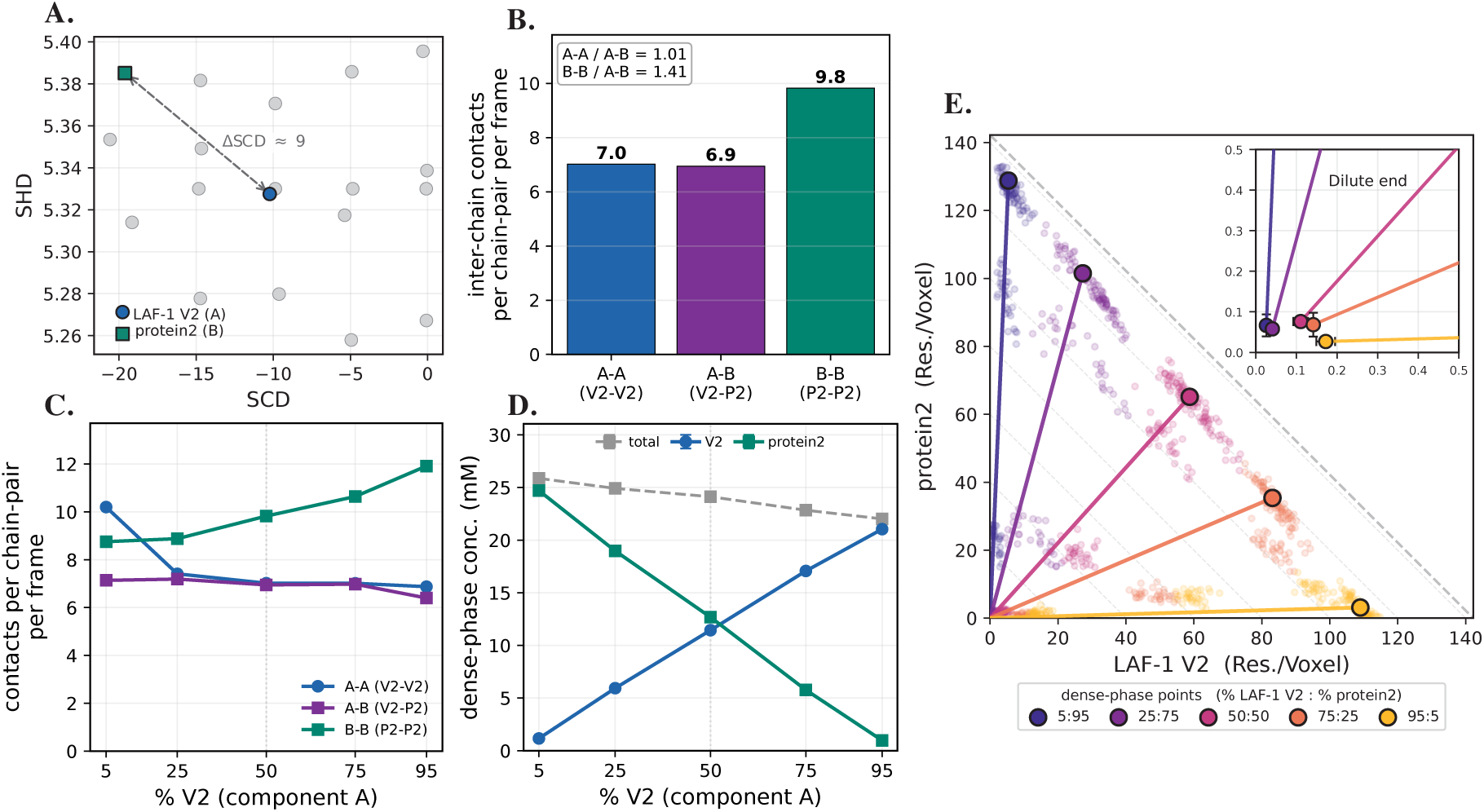
A strongly self-associating sequence and the reference protein form a single homogeneous condensate despite their different patterning. A) The mixed pair on the SCD–SHD map highlighting V2 and P2. B) Mean intermolecular contacts per chain-pair per frame in the equimolar mixture, resolved into the three block types. C) Contact strength versus composition. D) Dense-phase concentration versus composition. E) Phase diagram shown using the domain-decomposition, per-voxel method. All error bars show block-averaged SEM.

We do not find it to be so simple. In the 50:50 condensate the V2–V2 and V2–P2 contacts are nearly the same per chain-pair per frame (Figure 3B). However, P2–P2 interactions occur significantly more frequently, introducing an asymmetry into the internal structuring of the mixed condensate. This is unsurprising because P2 is the most strongly self-associating sequence, but what is surprising is that the enhanced homotypic interaction does not exclude V2 from the P2-rich network. In fact, it forms cross-contacts with V2 as frequently as V2 interacts with itself.

As the ratio of components is varied, the three contact types no longer move together (Figure 3C). The V2–P2 cross-contacts remain nearly constant at around 7 across the full composition range. In contrast, both self-contact types change with composition. P2-P2 contacts are the strongest in most cases. They increase moderately from about 8.8 to 11.9 as P2 becomes the minority component. This suggests that P2 forms more self-contacts as it becomes diluted. A similar trend was observed for the matched partner in Figure 2C, but it appeared only at the most dilute (5%) composition. Here, the increase occurs gradually across the composition series. Intriguingly, when V2 is dilute, V2–V2 contacts jump upward to about 10 instead of decreasing as they did in the V2-P9 mixture. To understand the higher V2–V2 contact value, we examined the density profiles SI Figure 5A. The few V2 chains are located within the P2-rich slab, but appear to be somewhat localized toward the interface, showing a small excluding effect via competition with the much stronger P2-P2 interactions. This enrichment toward the surface is apparent up to 50:50, but not in the two cases when V2 is the majority component. Thus, V2 and P2 remain co-condensed, but the component with weaker interactions seems to be partially shuffled toward the exterior of the condensate. Despite this apparent demixing, the demixing index remains at or below 0.05 at all compositions SI Figure 5B. Its largest value is 0.047 at the 50:50 composition. These low values indicate that the two proteins remain mixed throughout the system with only moderate drive toward demixing.

The phase diagram shows a similar trend as before(Figure 3D). The concentration of each protein in the dense phase changes with its fraction in the mixture. Neither protein is preferentially enriched or excluded relative to its amount in the input mixture. The total concentration changes only from about 26 mM at 5% V2 to 22 mM at 95% V2, and reflects a linear combination of the single-component densities of V2 and P2 condensates.

Intriguingly, in the voxel phase diagram (Figure 3E), we find the dense-phase ends of the tie lines fall along a single anti-diagonal band, showing much smaller fluctuations in total condensate density than in the previous case. The dilute-phase ends are clustered near the origin, below about 0.2 residues per voxel. Due to degree of uncertainty, we cannot ascertain whether the dilute coexistence curve is linear or concave. The overall phase diagram indicates a single mixed condensate whose composition follows the input mixture.

The supporting panels (SI Figure 6) are consistent with a shared, well-mixed condensate. The three residue–residue contact maps are all populated over the same charged blocks near the sequence ends, with the P2–P2 map the large number of contacts of the three (SI Figure 6A–C). The V2–P2 cross-map shows that residues from V2 and P2 make direct contacts with one another SI Figure 6A,B. This agrees with Figure 3B, where V2–P2 contacts occur nearly as often as V2–V2 contacts.

Inside the condensate, both proteins remain similar in size, with radii of gyration of about 35–37 Å across most compositions (SI Figure 6D). V2 remains near 36–37 Å at all compositions, while P2 becomes more compact at dilute concentrations (SI Figure 6D,E). This is somewhat surprising as the P2 chains are at the center of a condensate in these cases, rather than at the interface. It may reflect an incompatibility of V2-P2 interactions, making the primarily V2-comprising condensate a poor solvent for a single chain of P2.

The two proteins also diffuse at different rates within the condensate (SI Figure 6F). V2 remains nearly constant at approximately 3 × 10*^−^*^6^ cm^2^ s*^−^*^1^. In contrast, P2 diffuses more slowly at most concentrations, and then increases to slightly faster than V2 (3.7 × 10*^−^*^6^ cm^2^ s*^−^*^1^) at the dilute condition. Across most compositions, P2 diffuses more slowly than V2. This difference remains even though both proteins are mixed within the same condensate and experience a similar local density. This is consistent with our previous observation that a more charge-segregated sequence with lower SCD diffuses more slowly at matched density [30]. The results also demonstrate that the two components can retain different mobilities within the same mixed phase.

### Mixing with a weak self-associator that partly separates

We then consider the opposite case where the partner protein is much less charge segregated, and less prone to phase-separate on its own. We use P15. Its critical temperature is 283.52 K (SI Figure 1), which is below the 298 K of our simulations. It is comparably distant from V2 in SCD space, albeit in the opposite direction with 10 units greater SCD (Fig. 4A). Thus, its charges are distributed more evenly rather than clustered into blocks, having a similar SCD value to the well-mixed wild-type sequence. It also has the highest SHD in the library (Figure 4A). Among the sequences with well-mixed charges, it has relatively strong self-association. However, it does not phase separate on its own at 298 K. At this temperature, P15 can enter a condensate when it is mixed with a partner that phase separates on its own. Since V2 phase separates on its own, it may act as the scaffold and P15 as the client [43].

**Figure 4:**
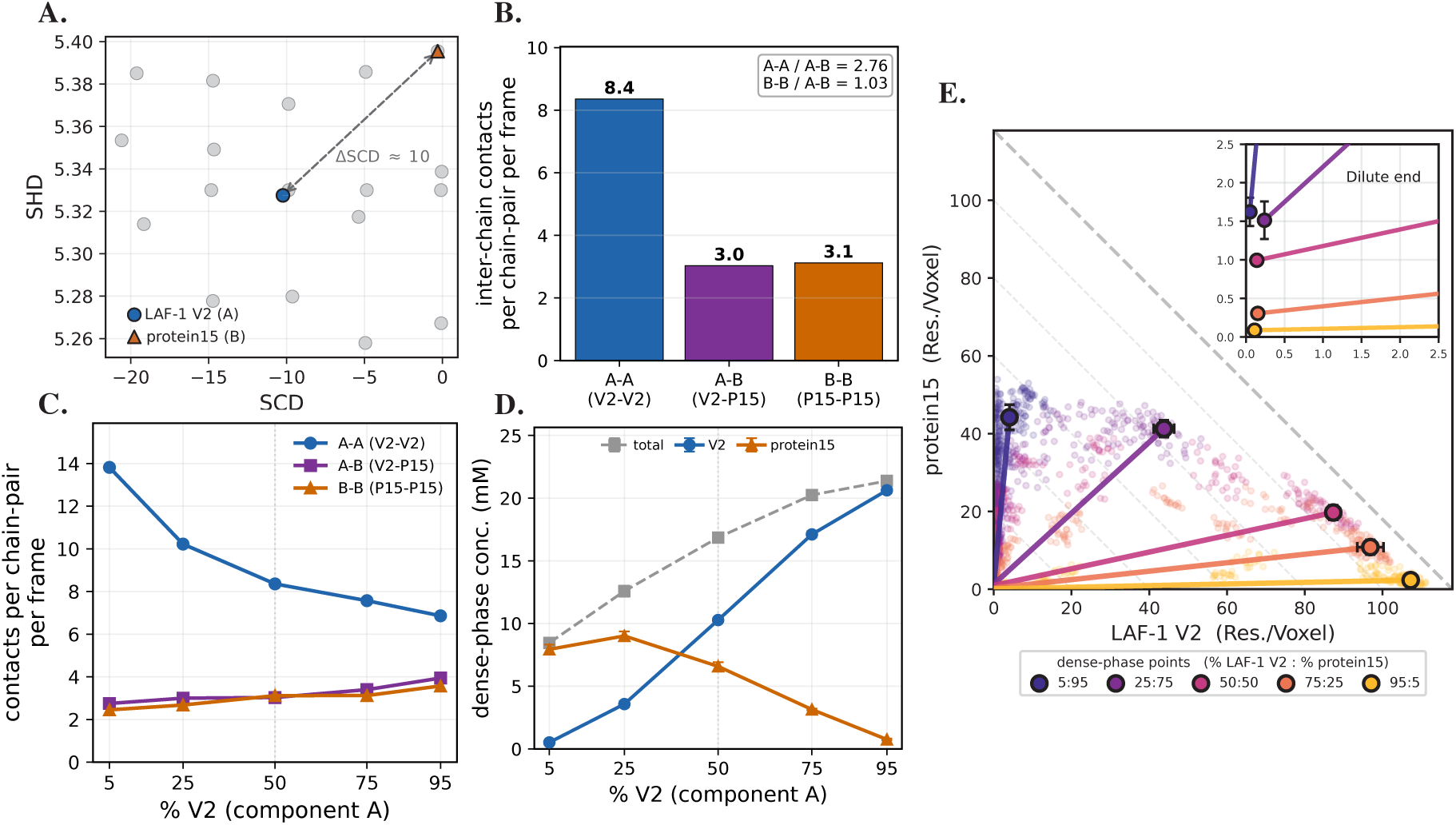
A weakly self-associating sequence and the reference protein form a scaffold–client system. A) The mixed pair on the SCD–SHD map highlighting V2 and P15. B) Mean intermolecular contacts per chain-pair per frame in the equimolar mixture, resolved into the three block types. C) Contact strength versus composition. D) Dense-phase concentration versus composition. E) Phase diagram shown using the domain-decomposition, per-voxel method. All error bars show block-averaged SEM.

As expected, in the 50:50 condensate the V2–V2 contacts lead at 8.4 per chain-pair per frame, well above the V2–P15 and P15–P15 counts of 3.0 and 3.1 (Figure 4B). V2 forms most of the interaction network, but P15 remains an active component of the condensate. Over the composition range the V2–V2 contact density falls from about 14 to 7 as V2 is diluted by P15, while the cross and B–B counts drift up, from about 2.5 to 4 (Figure 4C). So V2 always leads in number of interactions, but P15 stays involved at every mixing ratio. To see how the two proteins are arranged in space, we look at the density along the slab axis (SI Figure 7A). The condensate is not uniformly mixed. V2 forms a narrow, dense peak at the center of the slab. P15 forms a broader and less dense distribution that extends further on both sides. This pattern suggests a partial core–shell arrangement. V2 forms the protein-rich core, while P15 is more concentrated near the outer regions. The pattern is most visible at the 50:50 composition. The demixing index supports this pattern. It is close to zero at the composition extremes and increases to 0.34 at 50:50 (SI Figure 7B). This suggests partial separation between V2 and P15. The index is less informative at highly uneven compositions. At 5% and 25% V2, there is little V2 available to redistribute. The local voxel compositions therefore remain close to the overall ratio, and *D*_demix_ stays near zero. A low value at these compositions does not necessarily indicate uniform mixing. Instead, it shows that the index has limited sensitivity when one component is dilute. The density profiles provide a clearer view at these compositions. At 25% V2, they already show V2 gathering near the center of the slab. This feature is not captured by the demixing index. We therefore use the density profiles to interpret these systems.

The concentrations show a partial, though not complete, exclusion (Figure 4D). V2 builds the dense phase from about 0.5 to 21 mM as its fraction rises, while P15 is pushed down but never out. Its dense-phase concentration stays finite across the whole range, from about 8 mM at the P15-rich end down to about 0.7 mM at the V2-rich end. The total condensate density changes with composition, unlike in the previously-considered mixtures. For P15, the total dense-phase concentration increases as the V2 fraction increases. It rises from about 8.5 mM at 5% V2 to about 21 mM at 95% V2 (Figure 4D). We note that there is not a stable dense slab formed in the two cases with majority P15 protein as 298K is greater than the *T_c_* for P15.

The dense-phase density increases from about 50 residues per voxel at low V2 fractions to about 110 when V2 is the majority Figure 4E. This supports the scaffold–client picture [51]. At a 50:50 input composition, it contains about 87 V2 and 20 P15 residues per voxel, having a significant enrichment of V2 compared to P15.

We find that the dominant interactions in the V2–V2 contact map shows significantly more contacts compared to interactions involving P15 (SI Figure 8A-C). The two weaker contact maps also show different patterns. V2–P15 contacts are mainly found in the charged end blocks. These are the same regions V2 uses for its own contacts. In contrast, P15–P15 contacts are spread across the middle of the sequence. This matches P15’s more evenly distributed charges and centrally located hydrophobic residues.

The chain sizes are clearer once we look at which chain actually changes. P15 remains extended across the full composition range. Its *R_g_* stays near 35 Å, and its scaling exponent remains near 0.535 (SI Figure 8D,E), possibly since its *T_c_* is very close to 298K and it would thus behave similar to an ideal chain regardless of its presence in a condensate or in bulk. V2, however, shows a larger change. Its *R_g_* is about 28 Å at 5% V2 and increases to about 37 Å as V2 becomes the majority. Its scaling exponent also increases from about 0.52 to 0.55. This is most likely due to the disappearance of a stable slab in the most dilute condition. The two proteins also diffuse at different rates (SI Figure 8F). As V2 increasingly forms its own interaction network, its diffusivity decreases from approximately 8 × 10*^−^*^6^ to 3 × 10*^−^*^6^. However, P15 remains more mobile and shows little composition-dependent change, with diffusivities. We suspect part of the difference may reflect the location of P15 rather than its interactions alone.

This provides a contrasting case to the previous mixing pairs we considered. P15, unlike the P2, has little intrinsic tendency to form P15–P15 contacts which seems to limit its ability to build a persistent network within the dense phase. Moreover, its attraction to V2, is outcompeted by V2-V2 self-interactions which are stronger. So P15 is recruited only weakly into the V2-rich condensate. The two components start to separate, and the weaker P15 chains are pushed outward toward the interface. This is the partial-mixing case that sits opposite to the near-ideal mixing of P2. Importantly, this behavior could potentially be recapitulated by the V2-P2 case if the temperature were increased beyond the *T_c_* of V2, making V2 the client, and P2 the scaffold.

### (De)Mixing with a non-phase separating partner

We finally ask what happens when the partner has minimal self-association of its own and no leftover stickiness to make up for it using our least phase-separation-prone variant, P13. Like P15, it has a nearzero SCD (-0.08 against V2’s -10.23) and so barely self-associates, but unlike P15 it does not carry a high hydropathy pattern to compensate. Its SHD (5.27) is below V2’s (5.33), among the lower values in the library rather than the highest (Figure 5A), and its single-component critical temperature is 277.46 K (SI Figure 1). It is the partner with the least reason to join V2’s network, and that is what we see.

**Figure 5:**
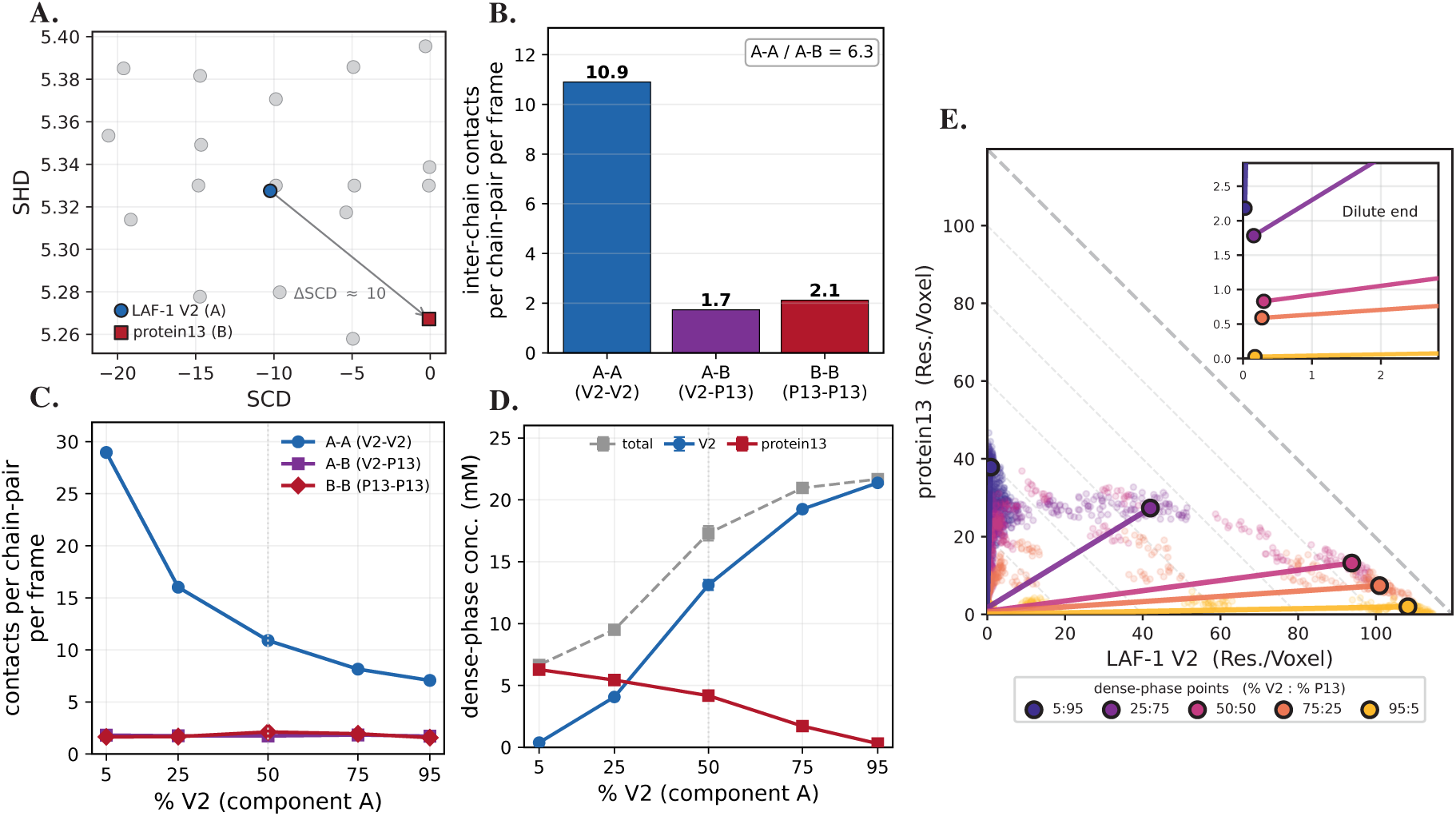
A non-self-associating sequence and the reference protein form a scaffold–client system. A) The mixed pair on the SCD–SHD map highlighting V2 and P13. B) Mean intermolecular contacts per chain-pair per frame in the equimolar mixture, resolved into the three block types. C) Contact strength versus composition. D) Dense-phase concentration versus composition. E) Phase diagram shown using the domain-decomposition, per-voxel method. All error bars show block-averaged SEM.

Splitting the contacts in the 50:50 mixture shows the most contacts by far for V2–V2, and many times lower contacts for pairs involving P13 (Figure 5B). SThis is the opposite of the balanced counts we observed previously for the similar and well-mixing case, and considerably more one-sided than the partial mixing case discussed in the previous section.

The same holds across the mixing range, and the shape of the V2–V2 curve is worth noting (Figure 5C). The V2–V2 count is highest when V2 is the minority, about 29 contacts per chain-pair at 5 % V2, and falls to about 7 as V2 becomes the majority. That looks backwards at first, it follows from the small number of V2 chains at the low-V2 end of the mixing range. The V2 chains in the minority will cluster together leading to a large V2–V2 contact count when normalized per pair (Figure 5C). Throughout the mixing range, both the V2–P13 and P13–P13 contact counts remain low and nearly constant, varying only from approximately 1.6 to 2.1. Thus, P13 does not become an active contributor to the condensate interaction network at any composition. The demixing index partially captures this behavior, though it is negligible at both ends of the mixing range, measuring 0.01 at 5% V2 and 0.005 at 95% V2 (SI Figure 9B). The index rises for intermediate compositions and peaks at 0.75 in the 50:50 mixture, consistent with strong demixing at equal proportions. For comparison the same index reached about 0.03 for the mixing pair in the near-ideal mixing case, and 0.34 for the partial demixing pair, so P13 sits clearly beyond both.

The dense-phase concentrations also show scaffold–client relationship, only forming a sufficiently large slab at the 50% or greater V2 (Figure 5D). V2 acts as the scaffold. Its dense-phase concentration increases from about 0.4 mM at 5% V2 to about 21 mM at 95% V2. P13 acts as the client, still entering the dense phase, but at a lower concentration than the expectation, given the concentration of V2 in the dense phase, i.e. having a significantly lower partition coefficient. The total dense-phase concentration is largely determined by the V2 concentration in most cases, and all cases involving a fully-formed slab.

The voxel phase diagram shows the same scaffold–client behavior (Figure 5E). At a 50:50 input composition, the dense phase contains about 94 V2 and 13 P13 residues per voxel, making it strongly enriched in V2. However, P13 remains present in the dense phase at all compositions, and generally has a higher concentration in the dense phase than outside, indicating that it is still preferentially incorporated rather than excluded. P13 also shows stronger separation from V2 than P15. At 50:50, the dense-phase V2:P13 ratio is about 7:1, compared with about 4:1 for V2:P15. The demixing index is also higher for P13 (*D*_demix_ = 0.74) than P15 (*D*_demix_ = 0.34). All this suggests that P13 separates more strongly from V2.

The supporting panels put this on a residue and single-chain footing (SI Figure 10). The three contact maps are on one shared color scale, and on that scale only the V2–V2 map carries real signal (SI Figure 10A–C). These contacts are concentrated in V2’s charged terminal blocks, around residues 0–30 and 140–165. The single-chain measurements follow the same pattern as the P15 mixture (SI Figure 10D–F). P13 remains similar in size and mobility across compositions. V2 shows the larger composition-dependent changes. This suggests that the separation is related to how the proteins associate within the condensate rather than differences in their individual chain properties.

This system shows scaffold–client behavior. V2 forms the dense phase, while P13 partitions into it more weakly and remains closer to the outer region. It does not show complete demixing within the dense phase, which would require two separate dense phases. What we see instead is P13 gathered near the surface or edge of the V2-rich dense region (75% and 95% V2) rather than mixed evenly through it (SI Figure 9A). At 5% V2 there is no clear condensate at all. P13 stays diffuse because it cannot phase separate on its own, and the small amount of V2 is mixed loosely into that P13-rich region.

### The mixing landscape across all partners

The previous sections considered multiple cases of co-phase separation, mixing and demixing, keeping LAF-1 V2 fixed as one partner and varying the second sequence. P9, whose charge pattern closely matches that of V2, mixed uniformly with V2; P2 was enriched more strongly in the condensate, while P15 and P13 were largely left out of V2 condensates. Those pairs make the point that mixing is set by how strongly the partner sticks to itself relative to how strongly it sticks to V2, but three pairs cannot say whether that reading holds across the library. Here we consider all 18 partners at the same five compositions in mixtures with V2 and look at the two quantities that measure mixing side by side.

We first calculated the *D_demix_* for all 18 partner sequences in each of the five stoichiometries with LAF-1 V2 (Figure 6A). For most of the library, *D_demix_* stays below the previously defined threshold for demixing of 0.2. There were four exceptions which break from that baseline, the wild-type (WT) LAF-1 RGG sequence, P13, P14, and P15. For these mixtures, the demixing index increases at intermediate compositions and reaches its largest values near the 50:50 ratio. Notably, these four partners also have the most well-mixed charge patterns, with SCD ≈ 0 (Figure 6A). They are also the only partners with single-component critical temperatures below the simulation temperature of 298 K (SI Figure 1). Thus we considered whether SCD might be a good predictor of demixing. At the composition extremes, the demixing index for these mixtures generally approaches zero.

**Figure 6:**
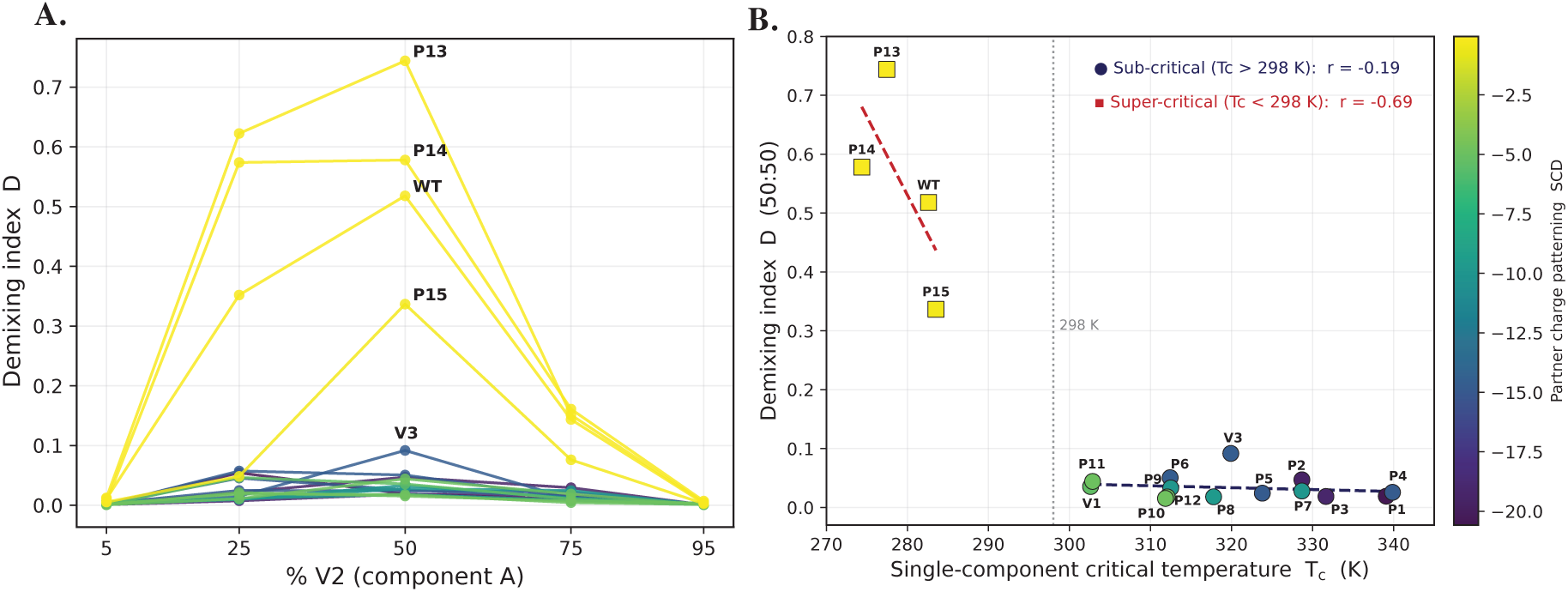
Collected mixing behavior of all V2-partner pairs. A) Demixing across all 18 partners mixed with LAF-1 V2 against composition for every partner. B) Demixing index, *D*_demix_, at a 50:50 composition of LAF-1 V2 and each partner at 298 K, plotted against the partner’s single-component critical temperature, *T_c_*.

Using the 50:50 case as the most prone to demixing, we plot the *D_demix_* against each partner’s single-component critical temperature. There does not appear to be a single clear relationship, aside from a rapid switch from mostly mixed to highly demixed, which occurs between sequences above and below their *T_c_*. We separated the sequence library into two groups (Figure 6B) to independent regression models, finding only weak correlation of *D_demix_* with *T_c_*. We did not find any improved correlation when considering charge patterning as a predictive property (SI Figure 11). Charge patterning follows the same trend because charge segregation promotes self-association and increases *T_c_*.

Generally, partners with *T_c_ >* 298 K phase separate on their own and remain well-mixed with V2. They have low *D_demix_* values across different charge patterns, only showing minimal predictive capacity of *T_c_* (Figure 5B). The four partners that show higher demixing have *T_c_ <* 298 K. These sequences do not phase separate on their own at 298 K, and their demixing index is more negatively correlated with *T_c_* (Figure 5B). However, given the small sample size, we do not suggest this as a general model without further testing.

Overall, the main difference in two-component mixing is whether the partner can phase separate on its own. Sequences that self-associate can more readily join a shared condensate with V2. Sequences with weaker self-association are less strongly recruited into the V2-rich dense phase. Patterning descriptors are useful for understanding single-chain behavior, but they do not fully determine the outcome of two-component mixtures. SCD and SHD influence chain size, self-association, and *T_c_*. As a result, sequences with the same composition can behave differently on their own. In mixtures with V2, a partner generally co-condenses when it can phase separate on its own. The sequences that do not phase separate at 298 K show greater separation from V2. However, this behavior also depends on mixture composition. At the composition extremes, the demixing index decreases because the minority component is too dilute to form a separate region. A large excess of one component also favors mixing of the minority component. Thus, a pair that separates near equal composition can appear more mixed when one component is in excess. The single-component phase behavior of each sequence can help predict its behavior with V2. However, the extent of demixing depends on both sequence properties and mixture composition.

## Methods

### Sequence library and patterning descriptors

The LAF-1 RGG variant library used throughout this study is the composition-matched sequence set developed in our previous work [30]. All 19 systems (the LAF-1 RGG wild type and 18 designed variants, hereafter denoted P1–P15 and V1–V3) have an identical length of *N* = 168 residues and amino-acid composition, differing only in the order of residues in the sequence.

Sequence patterning is quantified using two established descriptors. Sequence hydropathy decoration (SHD) is calculated from the scaled hydropathy value of each residue and its sequence separation [37, 38]:

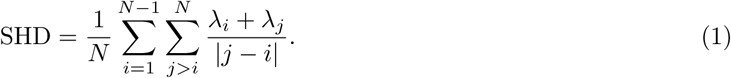

Here, *λ_i_* is the hydropathy of residue *i* on the Urry hydrophobicity scale [62]. Sequence charge decoration (SCD) is calculated from the formal charge and sequence separation of all residue pairs [37]:

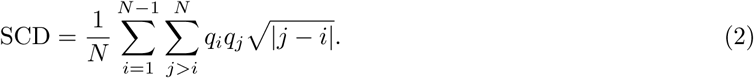

Here, *q_i_* is the formal charge of residue *i*. SCD becomes more negative as like charges are grouped into longer blocks. Because LAF-1 RGG carries almost no net charge, its wild-type sequence already lies near SCD ≈ 0. Reordering residues can therefore only increase charge segregation. The library spans SCD ≈ 0 to SCD ≈ −21.

### Coarse-grained model and simulation software

All molecular dynamics simulations used the HPS-Urry coarse-grained model [58, 62], in which each amino acid is represented by a single bead located at the residue’s center of mass. Simulations were performed with OpenMM [63] using a Langevin integrator with a time step of 10 fs (0.01 ps) and a per-residue friction coefficient of

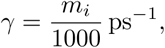

where *m_i_*is the molecular weight of residue *i*, following the mass-dependent friction scheme used in [58].

The total potential energy is the sum of one bonded term and two non-bonded terms, representing hydrophobic and electrostatic interactions:

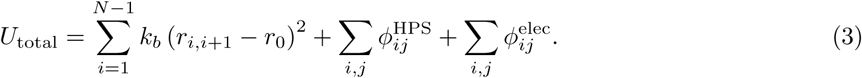

#### Bonded interactions

Adjacent residues within a chain are connected by a harmonic bond with equilibrium length *r*_0_ = 0.381 nm and spring constant *k_b_* = 8360 kJ mol*^−^*^1^ nm*^−^*^2^.

#### Hydrophobic interactions

Non-bonded hydrophobic contacts were modeled using the Ashbaugh– Hatch form of the Lennard-Jones potential [64]:

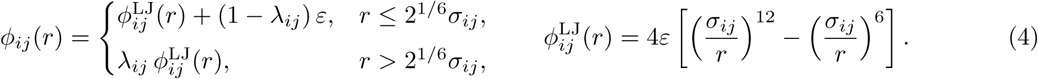

The energy scale was *ε* = 0.2 kcal mol*^−^*^1^ (0.837 kJ mol*^−^*^1^), the pairwise size was *σ_ij_* = (*σ_i_* +*σ_j_*)*/*2, and the pairwise stickiness was *λ_ij_* = (*λ_i_* + *λ_j_*)*/*2. Per-residue *λ_i_* values were obtained from the Urry hydrophobicity scale [62] and rescaled according to

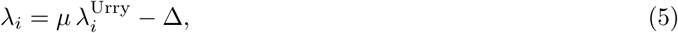

using the optimized HPS-Urry parameters *µ* = 1.0 and Δ = 0.08 [58]. Values of *λ_i_* below zero were set to zero. The hydrophobic potential was truncated at *r*_cut_ = 2.0 nm.

#### Electrostatic interactions

Charge–charge interactions were modeled using a Debye–Hückel potential [65], shifted to zero at the cutoff distance:

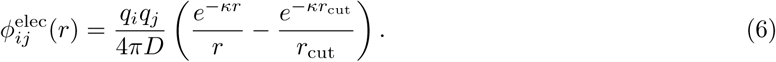

Here, *D* is the dielectric constant of water which we set to a constant value of 80. The inverse Debye screening length was set to *κ* = 1 nm*^−^*^1^, corresponding to a monovalent salt concentration of approximately 100 mM and consistent with previous HPS-Urry applications [58]. Histidine was assigned a charge of +0.5 *e* to represent partial protonation at neutral pH. The electrostatic potential was truncated and shifted at *r*_cut_ = 3.5 nm.

### Single-chain simulations

For each of the 19 LAF-1 RGG sequences, a single isolated chain was simulated at 298 K for 200 ns of production following 10 ns of equilibration.The trajectories were used to compute the mean single-chain radius of gyration, ⟨*R_g_*⟩ (Figure 1B), which decreases with increasing charge segregation.

### Single-component (multi-chain) slab simulations

For each of the 19 sequences, a slab-geometry simulation containing 100 chains was performed at 298 K in a periodic simulation box of 15 × 15 × 280 nm^3^, following the slab methodology established previously [30, 66]. Each system was equilibrated for 20 ns with a weak harmonic restraint that confined chains to the center of the simulation box, followed by 1 *µ*s of unrestrained production. A total of 5000 frames were saved at intervals of 0.2 ns.

These simulations were used to determine the dense- and dilute-phase concentrations at 298 K (Figure 1C,D) and multi-chain properties (Figure 1E,F; see below). Critical temperatures, *T_c_*, for these sequences (SI Figure 1) were obtained previously from multi-temperature slab simulations fit to coexistence curves assuming three-dimensional Ising universality, as described previously [30].

### Two-component slab simulations

Mixtures of LAF-1 V2 (component A, the reference sequence) with each of the other 18 library members (component B) were simulated at five compositions with a fixed total of 100 chains. Because all sequences have the same length, chains of both components were combined directly in a single slab-geometry simulation without modifying the single-component protocol. All two-component simulations were performed at 298 K in a periodic simulation box of 15 × 15 × 280 nm^3^.

Each system was equilibrated for 20 ns under a weak harmonic restraint,

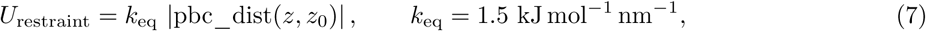

which confined all beads toward the center of the simulation box, where *z*_0_ = *L_z_/*2, under the minimum-image convention. This was followed by 1 *µ*s of unrestrained production without the restraint, saving 5000 frames at intervals of 0.2 ns.

### Domain-decomposition analysis of dense- and dilute-phase composition

Dense- and dilute-phase concentrations, as well as the condensate composition in two-component mixtures, were determined using the domain-decomposition method (DDM) of Morton and Vácha [57]. The simulation box was partitioned into cubic voxels with edge lengths of 35 Å.

The number of residues from each component was accumulated in each voxel and averaged over the production trajectory. For each component, the dilute-phase concentration was defined as the mode of the near-zero peak in the per-voxel residue-density distribution, estimated using kernel density estimation (KDE). The dense-phase concentration was obtained from the KDE restricted to voxels within the slab, defined as voxels with total residue density greater than the valley separating the dilute- and dense-phase populations in the total-density distribution.

For two-component phase diagrams, the plotted dense-phase composition of each species was calculated as the mean over voxels in the condensed core, defined as those with a total density exceeding the dense-phase peak of the total-density distribution. This quantity reflects the joint composition of the coexisting dense phase rather than the density mode of either individual component. Voxel residue counts were converted to molar concentrations using the voxel volume and the number of residues per chain.

### Demixing index

To quantify whether the two components of a mixture are spatially mixed or segregated within the condensate, we calculated the demixing index, *D*_demix_, following the method of Morton and Vácha [57]. For each voxel *i*, the pair of component densities, (*d*_1*,i*_*, d*_2*,i*_), was projected onto the global composition direction and onto the direction perpendicular to it. This procedure yields two variances: the spread along the composition axis, 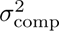, and the spread perpendicular to that axis, 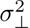. The demixing index compares these two variances:

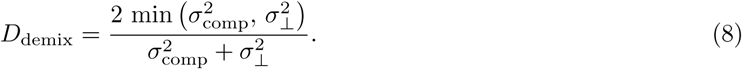

Here, *D*_demix_ = 0 when the voxel-to-voxel composition scatter is isotropic, corresponding to uniform mixing. In contrast, *D*_demix_ → 1 when nearly all of the scatter lies along the composition axis, corresponding to spatially distinct, compositionally pure regions. We classify mixtures with *D*_demix_ *<* 0.2 as well-mixed.

### Chain dimensions, internal scaling, and translational diffusivity

For chains within the slab, including both single-component and two-component simulations, the mean radius of gyration, ⟨*R_g_*⟩, was computed for each chain and then averaged over all chains. The internal scaling exponent, *ν*, was obtained by calculating the mean spatial distance, ⟨*r*(*s*)⟩, between residue pairs separated by a sequence distance *s*, for values of *s* up to half the chain length. The resulting distances were fit to

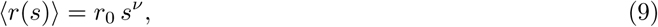

by linear regression in log–log space.

Translational diffusivity was computed from the chain center of mass using the continuous (never periodically wrapped) production trajectory. At every frame, each chain was reconstructed as a whole across the periodic boundary using minimum-image bond reconstruction before its center of mass was calculated. This avoids the spurious bounded or saturating displacements that result when mean-squared displacements are computed from box-wrapped coordinates.

For isolated single chains, the three-dimensional mean-squared displacement (MSD) of the center of mass was computed and fit by linear regression to the Einstein relation over an early, well-sampled lag-time window spanning 1%–10% of the trajectory length:

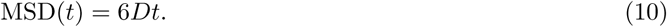

For chains within the dense phase of a slab, motion along the interface normal, *z*, is bounded by the condensate boundary and is therefore not representative of bulk translational mobility. We therefore used the in-plane (lateral, *xy*) displacement of the chain center of mass and fit the two-dimensional Einstein relation by linear regression over the same early lag-time window:

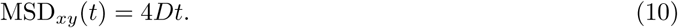

Thus, the fitting procedure is identical for free single chains and chains in the slab dense phase, except that the former uses unrestricted three-dimensional displacements, whereas the latter uses only in-plane, two-dimensional displacements in the *x*–*y* plane.

## Data and code availability

The data and code required to reproduce all figures in the main text and Supporting Information are available on GitHub at Dignon-Lab/Mixing_or_demixing_paper_figures. The repository is organized into one folder per figure, each containing a Python script and the property tables, contact maps, and DDM voxel exports used to generate that figure.

## Conclusion

We asked whether rearranging the same sequence residues can change the phase behavior of a disordered protein. We also examined whether these differences remain when two proteins are mixed. Our designed library of shuffled variants of LAF-1 RGG is well suited for this question. Each variant has the same length and composition but a different residue arrangement. We used a coarse-grained model and a library of composition-matched sequences. Each variant was tested with the same reference partner at a single temperature. This controlled setup allowed us to isolate the effect of sequence patterning from single-chain behavior to single-component and two-component condensates. On their own, the variants show that charge patterning, measured by SCD, strongly influences phase behavior. Sequences with segregated charges are more compact as single chains and form denser condensates at lower saturation concentrations than sequences with well-mixed charges.

We asked if these differences in phase separation propensity would persist within mixed condensates of multiple components, or whether the degree of mixing could be influenced by differences between sequences. Whether a partner mixes with V2 depends primarily on whether it can phase separate on its own at 298 K, rather than on how different its charge pattern is from V2. For example, P2 has a highly segregated charge pattern and differs from V2 by about nine SCD units. However, it self-associates strongly, with *T_c_* = 329 K and forms one mixed condensate with V2. P15 has well-mixed charges and differs from V2 in the opposite direction. It does not phase separate on its own at 298 K, with *T_c_* = 284 K. Instead, it shows partial separation, with a V2-rich core and a P15-enriched shell.

Across the library, demixing occurred only when one partner was above their single-component critical temperatures at 298K and therefore could not phase-separate on its own. Differences in charge or hydropathy patterning alone were not sufficient to cause demixing once the full interaction network of a crowded, multichain condensate was considered. Mixing also altered each protein’s internal behavior. The near-ideal chain scaling and matched dynamics observed in single-component condensates were no longer guaranteed, and proteins could retain distinct sizes, scaling behaviors, and mobilities even within the same dense phase. Because cellular condensates contain many components, predicting which proteins mix remains challenging. Our results suggest that a protein’s ability to phase-separate on its own may be a useful first predictor of whether it joins or is excluded from a condensate, reducing the need to compare every possible sequence pair. This principle could help explain cellular partitioning, assess disease mutations that alter self-association, and guide synthetic-condensate design. Extending it to larger interaction networks and real proteomes could turn descriptors such as SCD and SHD into predictors of condensate composition.

## Supporting information

### SI Figures

**Figure 1:**
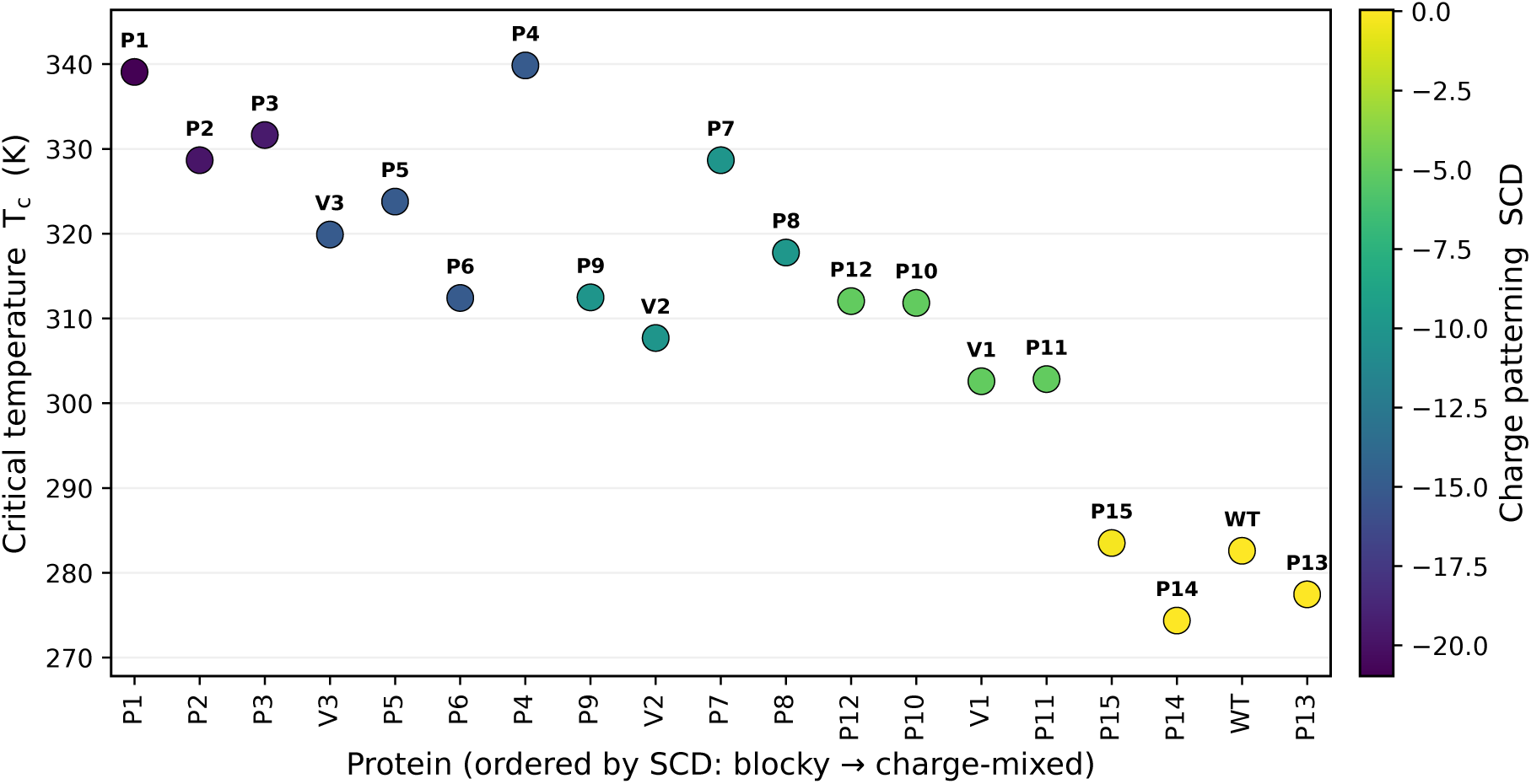
Single-component critical temperature increases with charge segregation. Critical temperature *T_c_* of each of the 19 LAF-1 RGG sequences from single-component slab simulations vs plotted per sequence.

**Figure 2:**
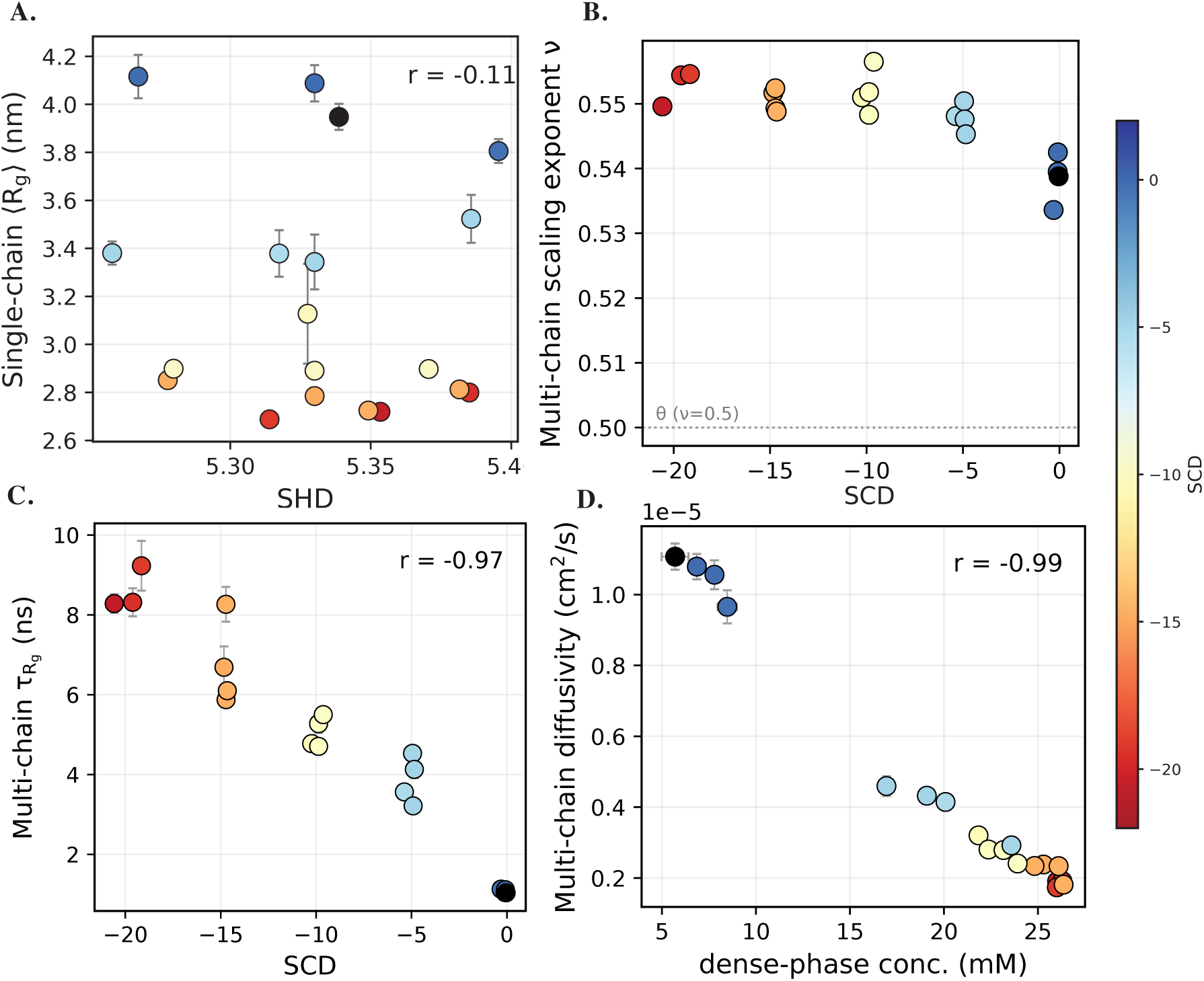
Supporting single-chain and single-component characterisation. (A) Single-chain *<R_g_>* versus SHD. (B) Multi-chain scaling exponent *ν* versus SCD. (C) Chain relaxation time *τ_Rg_* versus SCD. (D) Chain diffusivity D versus dense-phase concentration. Marker colour encodes SCD as in Figure 1. All error bars show block-averaged SEM.

**Figure 3:**
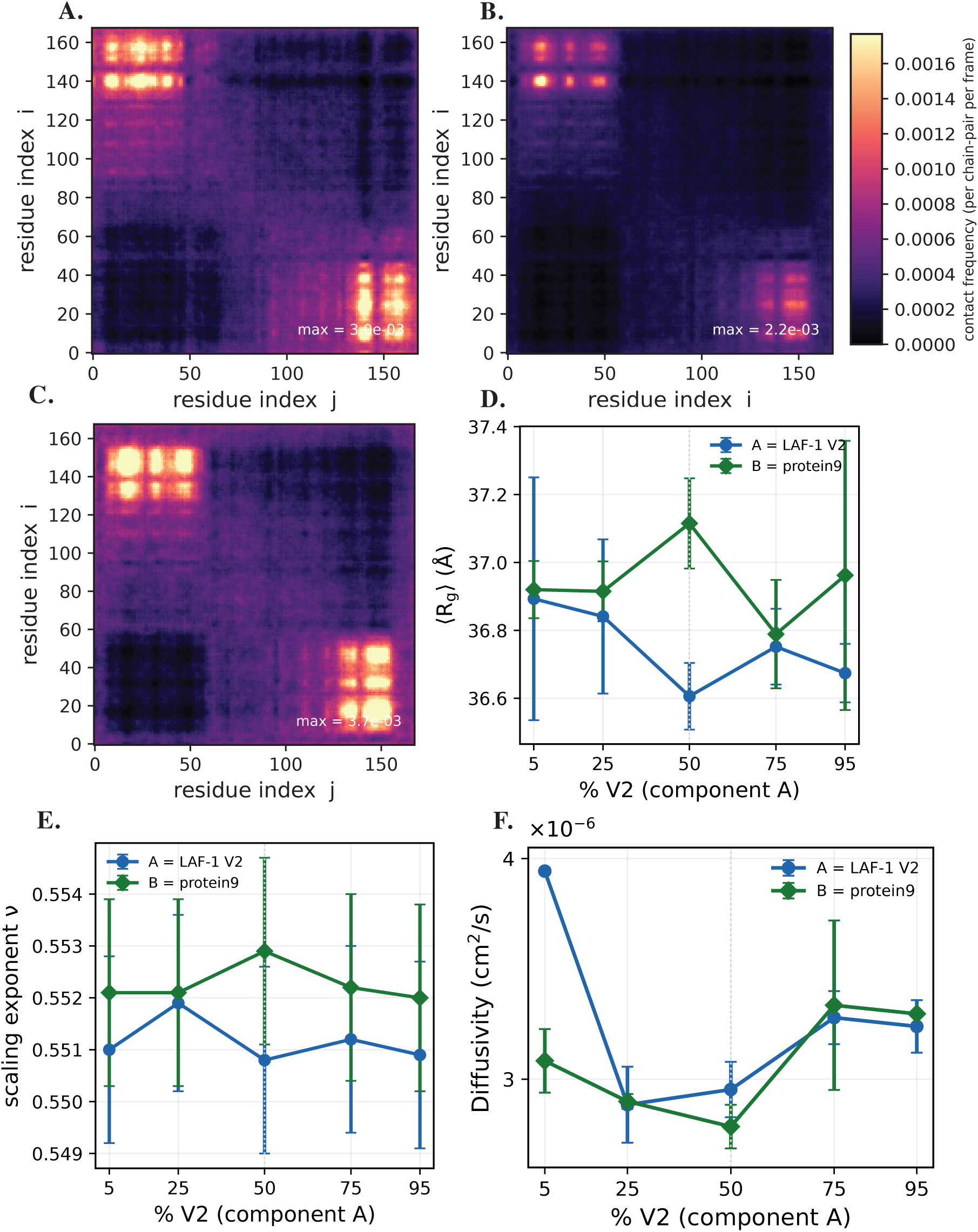
Ideal-mixing characterisation for LAF-1 V2 + protein9. (A–C) Intermolecular residue–residue contact maps for the A–A, A–B and B–B blocks on a shared colour scale. (A) A–A, V2 with V2. (B) A–B, V2 on the vertical axis (residue i) and protein9 on the horizontal axis (residue j). (C) B–B, protein9 with protein9. (D) Multi-chain single-chain *<R_g_>* versus composition. (E) Multi-chain scaling exponent *ν* versus composition. (F) Chain translational diffusivity D versus composition (log axis). All error bars show block-averaged SEM.

**Figure 4:**
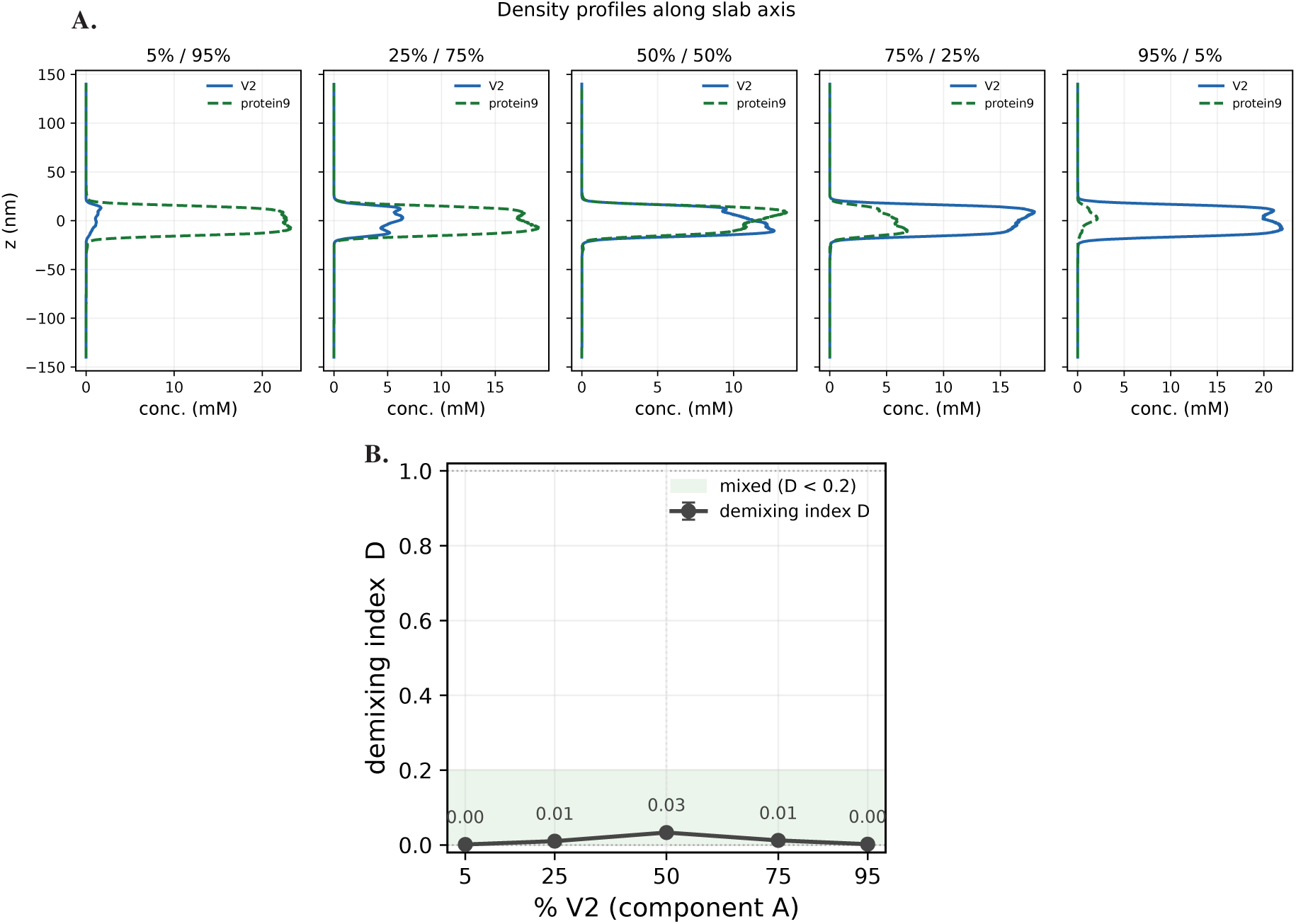
LAF-1 V2 and protein9 occupy the same dense slab and stay mixed at every composition (298 K). (A) Time-averaged density profiles (concentration in mM) of V2 (solid blue) and protein9 (dashed green) along the slab axis z, for the five mixing ratios. (B) Demixing index D versus composition (% V2).

**Figure 5:**
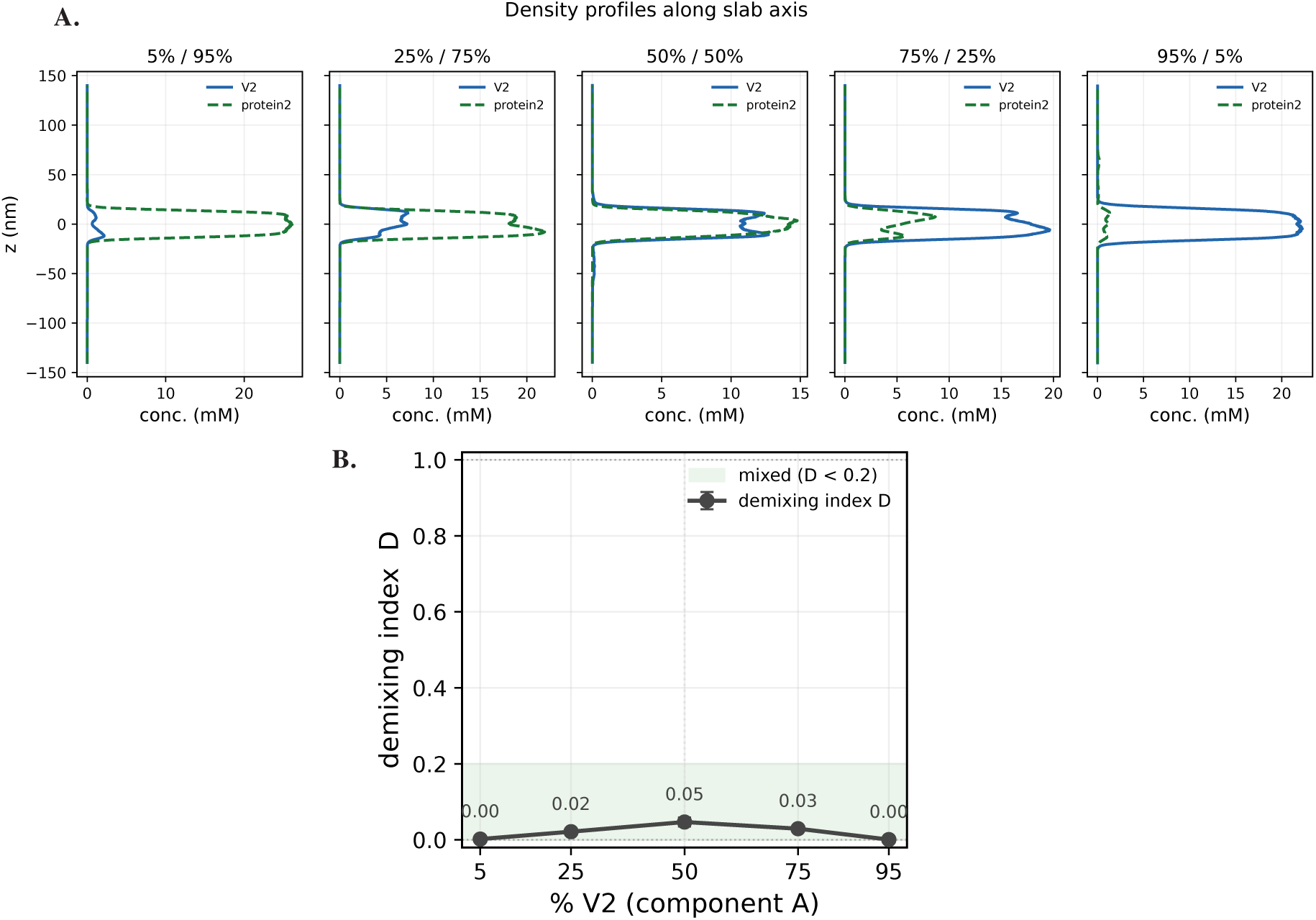
Density profiles and demixing index for LAF-1 V2 + protein2 across composition. (A) One-dimensional concentration profiles along the slab long axis, at each mixing ratio, with V2 (solid blue) and protein2 (dashed green) shown together. (B) Demixing index D versus composition (error bars are block-averaged SEM).

**Figure 6:**
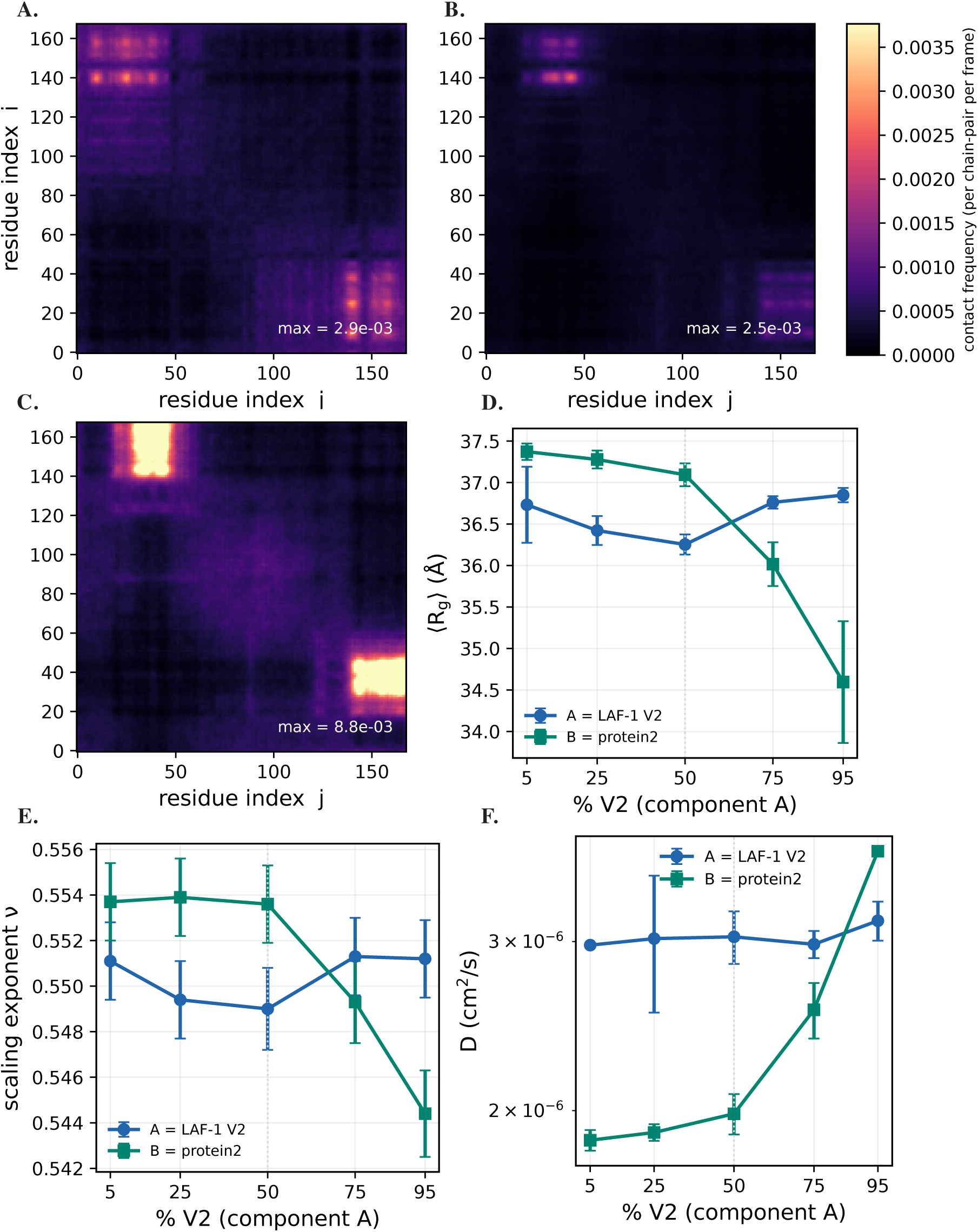
A strong self-associator characterisation for LAF-1 V2 + protein2. (A–C) Residue–residue contact maps for the A–A, A–B and B–B pairs on a shared colour scale. (A) A–A, V2 with V2. (B) A–B, V2 on the vertical axis (residue i) and protein2 on the horizontal axis (residue j). (C) B–B, protein2 with protein2. (D) Multi-chian single-chain *<R_g_>* versus composition. (E) Multi-chain scaling exponent *ν* versus composition. (F) Chain translational diffusivity D versus composition (log axis). All error bars show block-averaged SEM.

**Figure 7:**
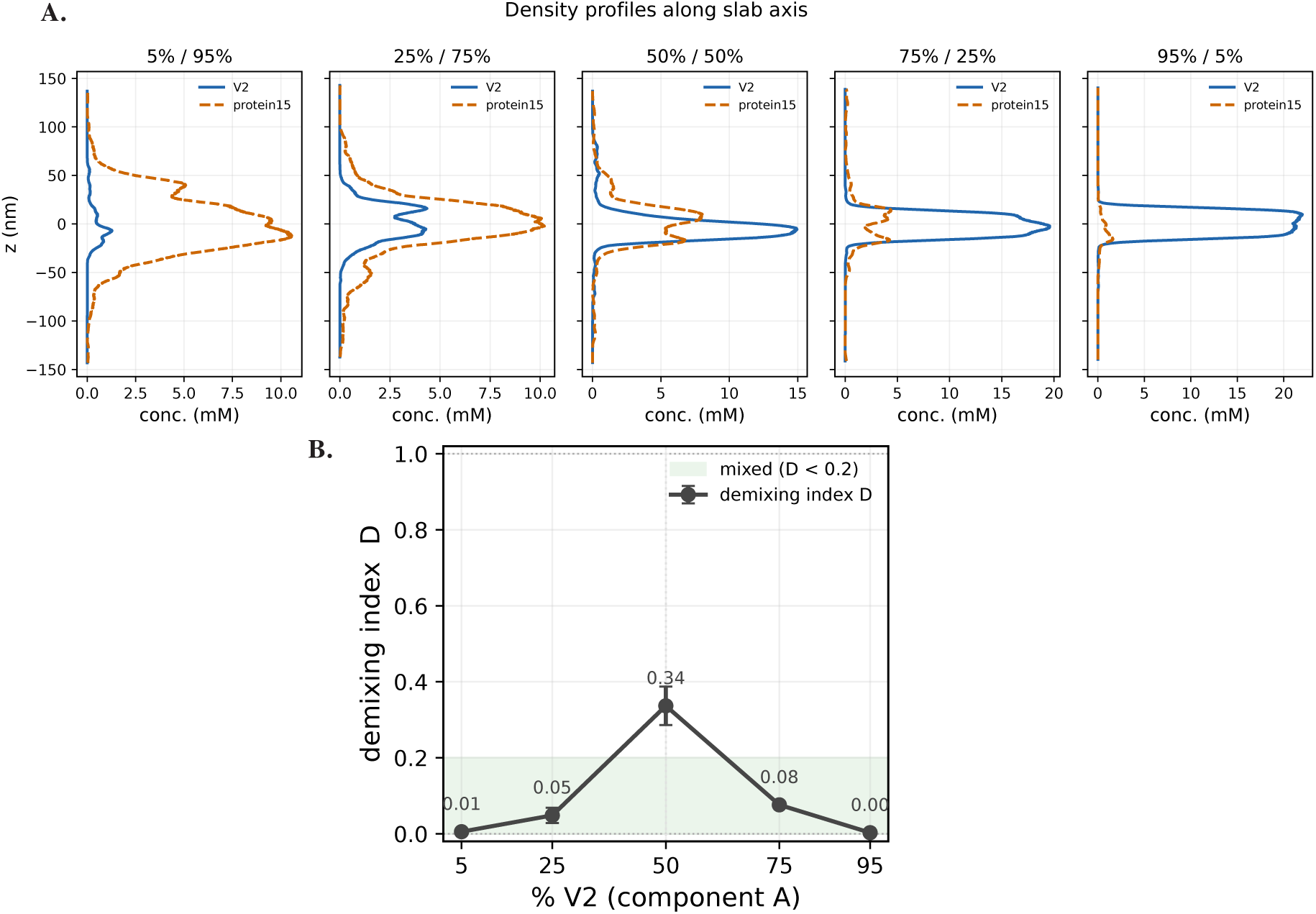
Density profiles and demixing index for LAF-1 V2 + protein15 across composition. (A) Concentration profiles along the slab axis at each mixing ratio, with V2 (solid blue) and protein15 (dashed orange). (B) Demixing index D versus composition (block-averaged SEM).

**Figure 8:**
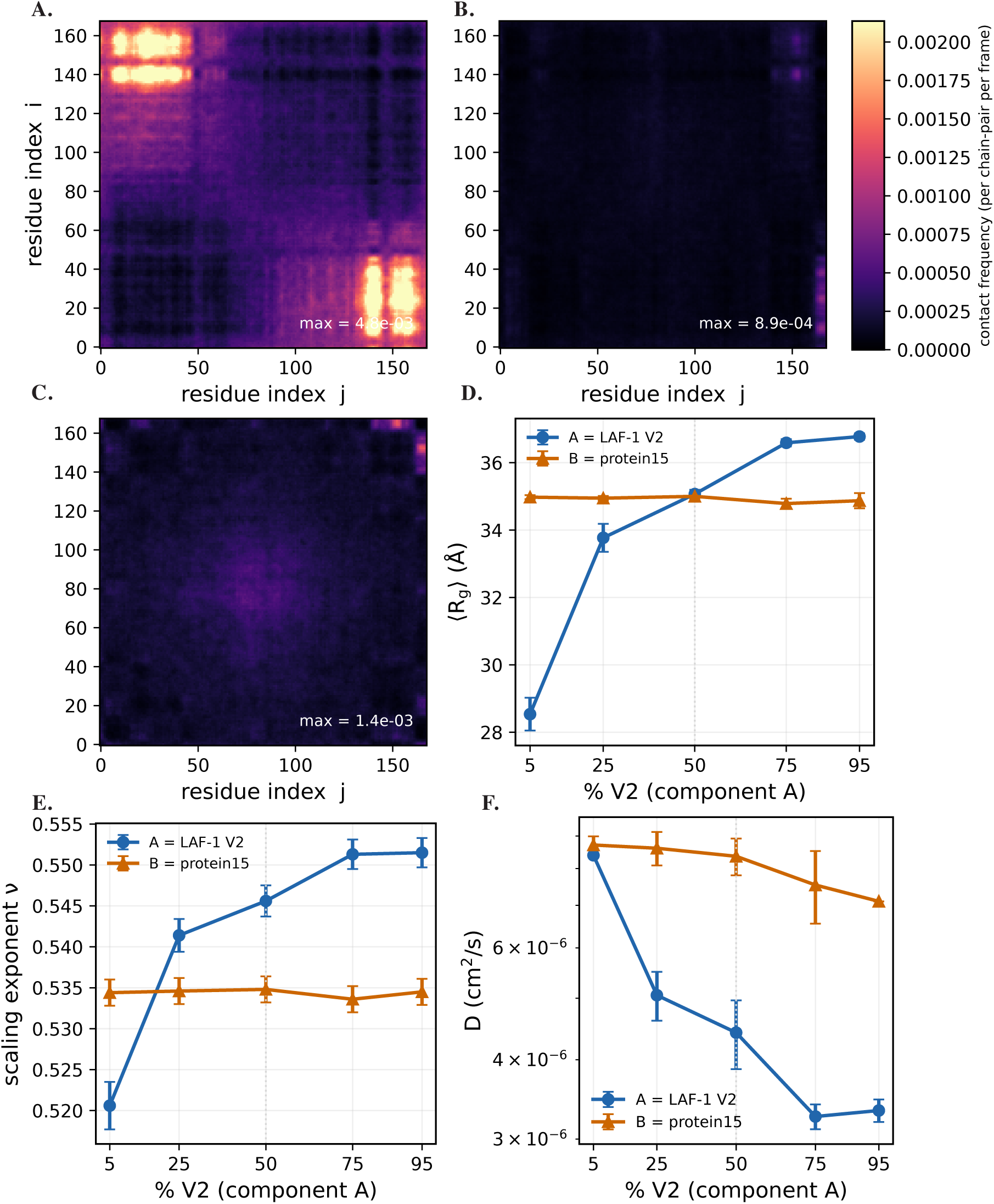
A weak self-associator partly separates from LAF-1 V2 (LAF-1 V2 + protein15). (A–C) Residue–residue contact maps for the A–A, A–B and B–B pairs on a shared colour scale. (A) A–A, V2 with V2. (B) A–B, V2 on the vertical axis (residue i) and protein15 on the horizontal axis (residue j). (C) B–B, protein15 with protein15. (D) Multi-chian single-chain *<R_g_>* versus composition. (E) Multi-chain scaling exponent *ν* versus composition. (F) Chain translational diffusivity D versus composition (log axis). All error bars show block-averaged SEM.

**Figure 9:**
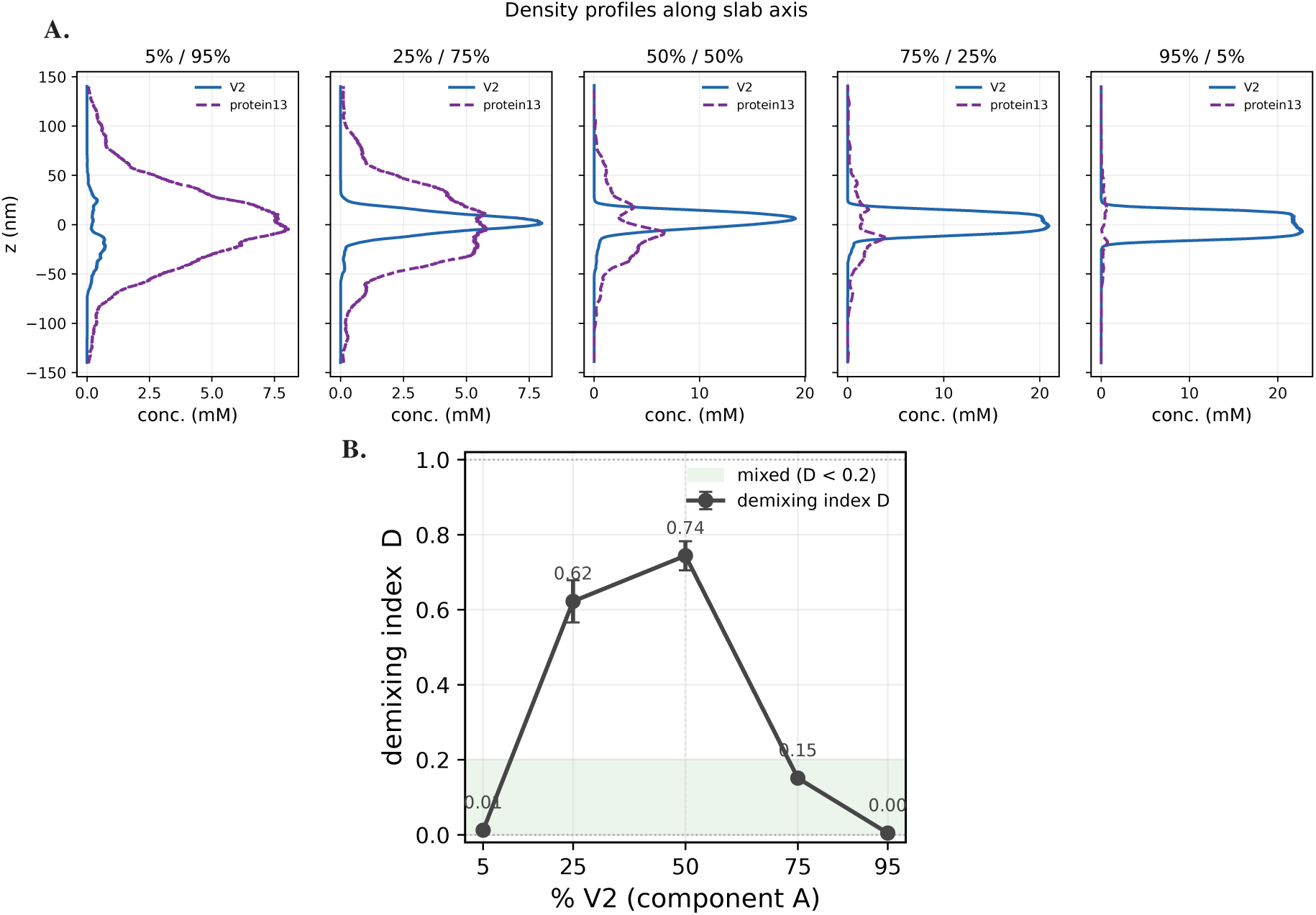
Slab density profiles and the demixing index for LAF-1 V2 + protein13 across composition (298 K). (A) Time-averaged concentration profiles along the long slab axis z for the two components, LAF-1 V2 (blue, solid) and protein13 (purple, dashed) at the five mixing ratios. (B) Demixing index D versus composition, from the DDM voxel analysis (35 Ä grid). All error bars show block-averaged SEM.

**Figure 10:**
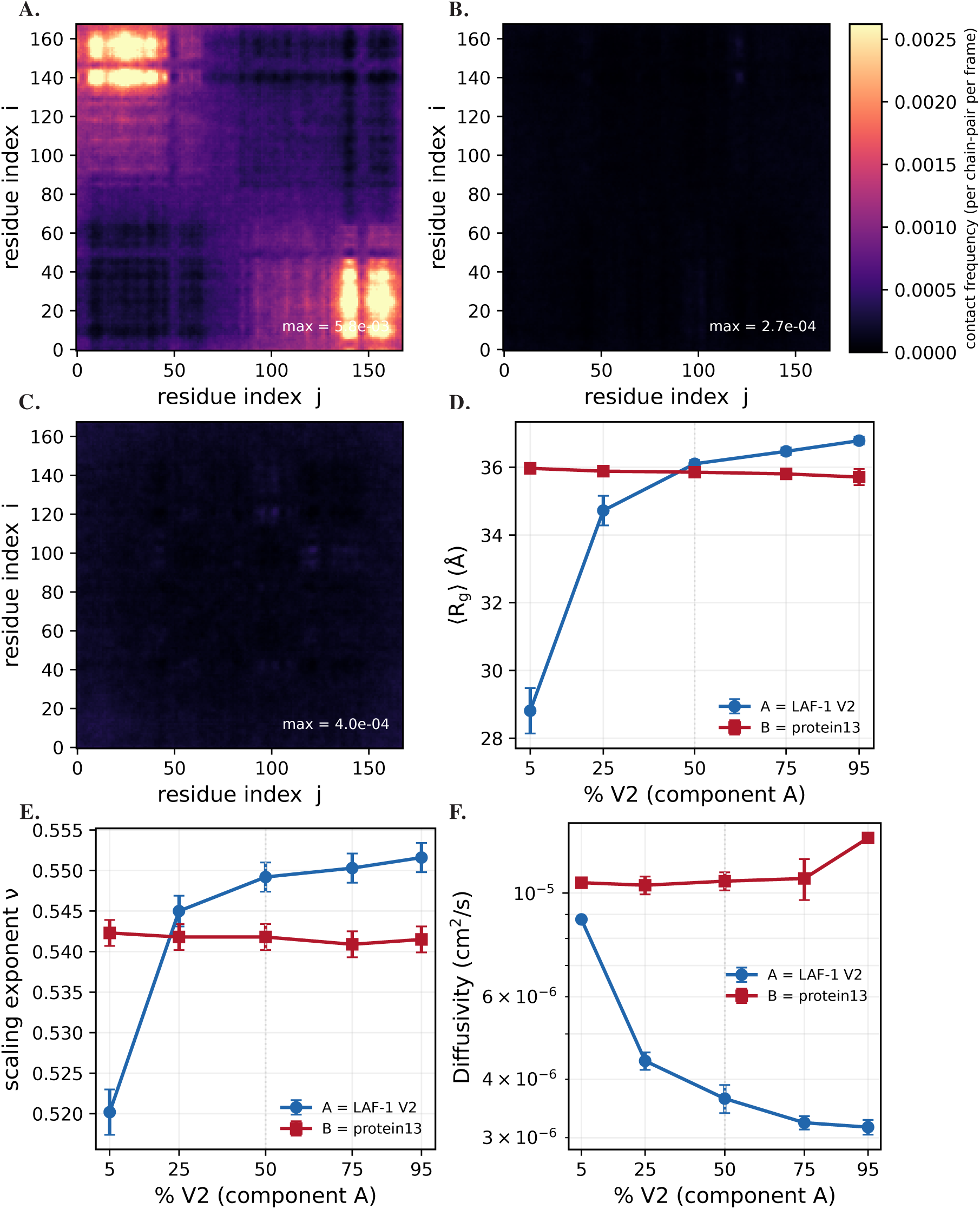
Demixing characterisation for LAF-1 V2 + protein13. (A–C) Residue–residue contact maps for the A–A, A–B and B–B pairs on a shared colour scale. (A) A–A, V2 with V2. (B) A–B, V2 on the vertical axis (residue i) and protein13 on the horizontal axis (residue j). (C) B–B, protein13 with protein13. (D) Multi-chian single-chain *<R_g_>* versus composition. (E) Multi-chain scaling exponent *ν* versus composition. (F) Chain translational diffusivity D versus composition (log axis). All error bars show block-averaged SEM.

**Figure 11:**
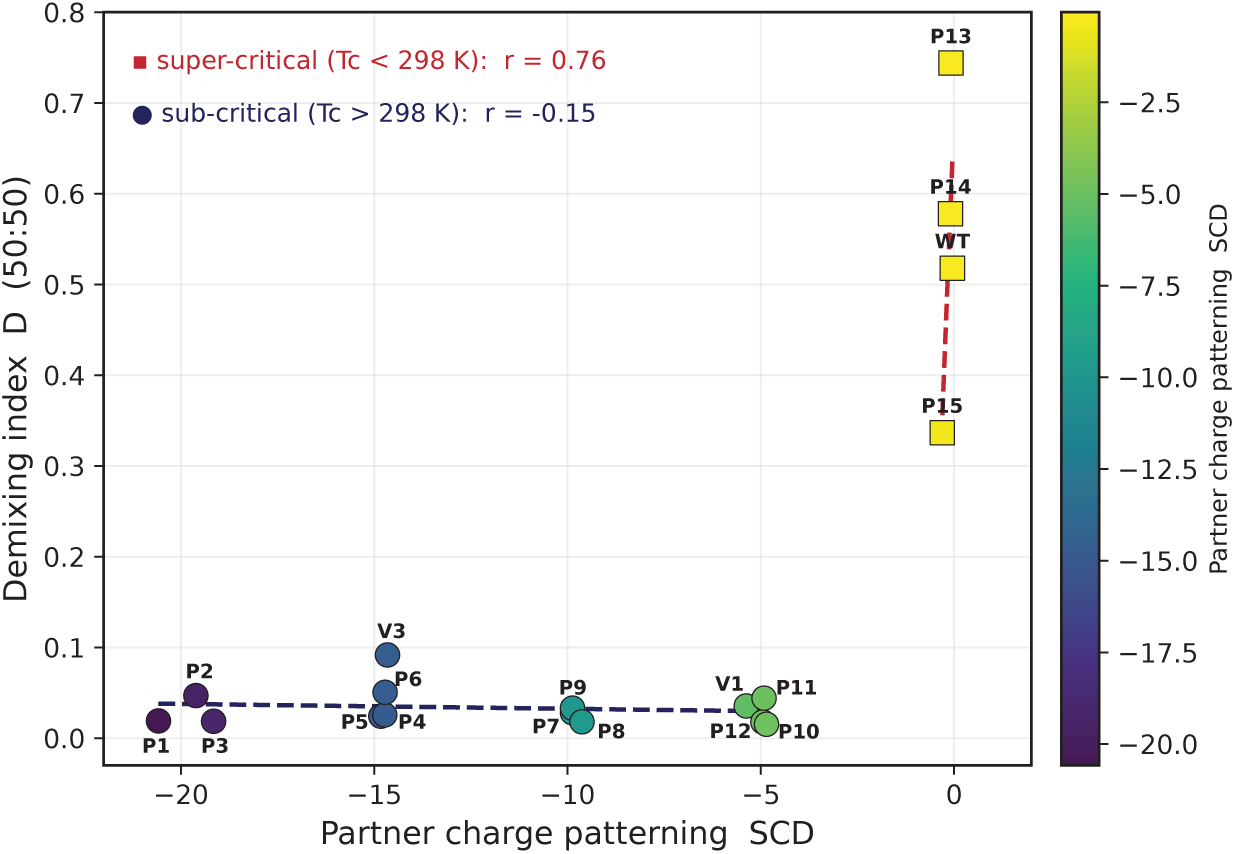
Demixing index versus charge patterning. Demixing index, D, at a 50:50 composition of LAF-1 V2 and each partner at 298 K, plotted against the partner’s SCD.

### SI Tables

**Table S1:**
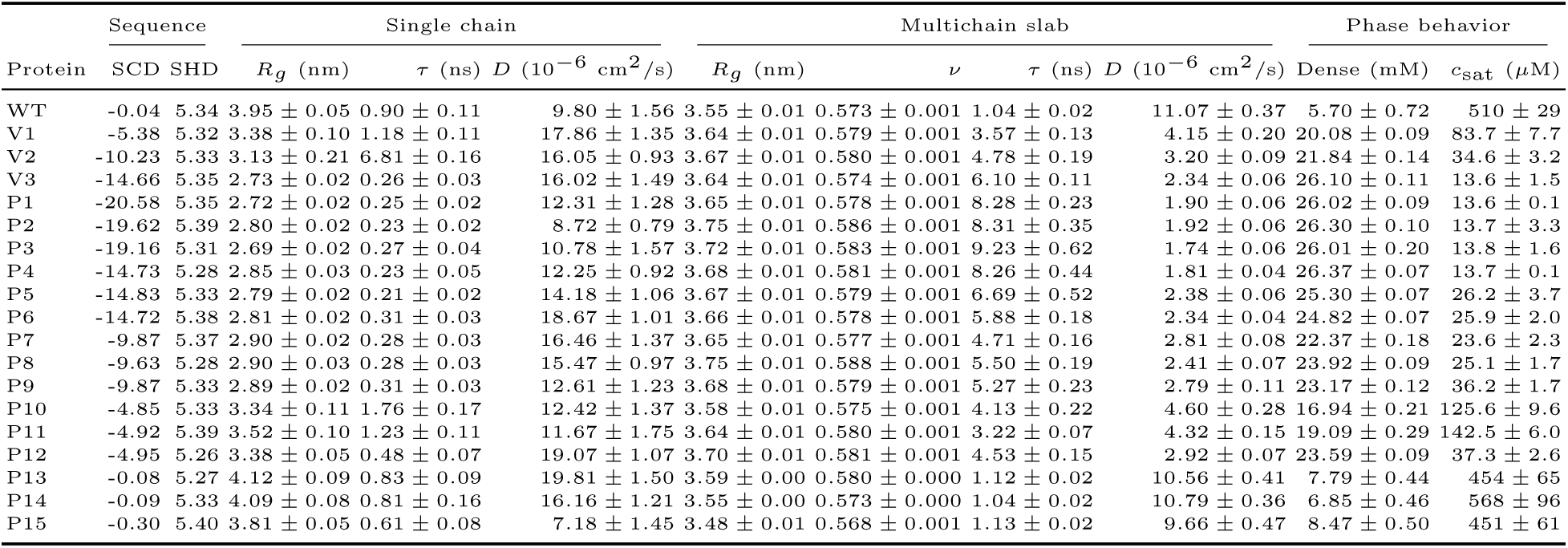
Single-chain and single-component (multi-chain) properties of the LAF-1 RGG sequence library (298 K). Values are mean *±* SEM. Diffusivities are reported in units of 10*^−^*^6^ cm^2^/s.

## Acknowledgments

G.L.D acknowledges support from NIH Award R35GM150589 and start-up funds from Rutgers University, and computing resources through NSF ACCESS DISCOVER project CHM250012. We thank José Villegas and Ben Schuster for useful discussions.

## References

(1) Banani, S. F.; Lee, H. O.; Hyman, A. A.; Rosen, M. K. Nature reviews Molecular cell biology 2017, 18, 285–298.

(2) Shin, Y.; Brangwynne, C. P. Science 2017, 357, eaaf4382.

(3) Brangwynne, C. P.; Eckmann, C. R.; Courson, D. S.; Rybarska, A.; Hoege, C.; Gharakhani, J.; Jülicher, F.; Hyman, A. A. Science 2009, 324, 1729–1732.

(4) Molliex, A.; Temirov, J.; Lee, J.; Coughlin, M.; Kanagaraj, A. P.; Kim, H. J.; Mittag, T.; Taylor, J. P. Cell 2015, 163, 123–133.

(5) Feric, M.; Vaidya, N.; Harmon, T. S.; Mitrea, D. M.; Zhu, L.; Richardson, T. M.; Kriwacki, R. W.; Pappu, R. V.; Brangwynne, C. P. Cell 2016, 165, 1686–1697.

(6) Larson, A. G.; Elnatan, D.; Keenen, M. M.; Trnka, M. J.; Johnston, J. B.; Burlingame, A. L.; Agard, D. A.; Redding, S.; Narlikar, G. J. Nature 2017, 547, 236–240.

(7) Rostam, N.; Ghosh, S.; Chow, C. F. W.; Hadarovich, A.; Landerer, C.; Ghosh, R.; Moon, H.; Hersemann, L.; Mitrea, D. M.; Klein, I. A., et al. Nature Methods 2023, 20, 673–676.

(8) Peeples, W.; Rosen, M. K. Nature chemical biology 2021, 17, 693–702.

(9) Küffner, A. M.; Linsenmeier, M.; Grigolato, F.; Prodan, M.; Zuccarini, R.; Capasso Palmiero, U.; Faltova, L.; Arosio, P. Chemical Science 2021, 12, 4373–4382.

(10) Riback, J. A.; Katanski, C. D.; Kear-Scott, J. L.; Pilipenko, E. V.; Rojek, A. E.; Sosnick, T. R.; Drummond, D. A. Cell 2017, 168, 1028–1040.

(11) Aierken, D.; Aland, S.; Bo, S.; Boeynaems, S.; Cai, D.; Carra, S.; Case, L. B.; Chan, H. S.; Espinosa, J. R.; GrandPre, T. K., et al. PRX Life 2026, 4, 032501.

(12) Alberti, S.; Gladfelter, A.; Mittag, T. Cell 2019, 176, 419–434.

(13) Patel, A.; Lee, H. O.; Jawerth, L.; Maharana, S.; Jahnel, M.; Hein, M. Y.; Stoynov, S.; Mahamid, J.; Saha, S.; Franzmann, T. M., et al. Cell 2015, 162, 1066–1077.

(14) Aguzzi, A.; Altmeyer, M. Trends in cell biology 2016, 26, 547–558.

(15) Li, X.; Wang, H.; Yao, J.; Han, B.; Zhao, X.; Jiang, Y.; Chen, H.; Yang, Y.; Hou, H.; Wang, L. Theranostics 2026, 16, 2684.

(16) Jiang, L.; Kang, Y. Biochimica et Biophysica Acta (BBA)-Reviews on Cancer 2025, 1880, 189245.

(17) Bah, A.; Forman-Kay, J. D. Journal of Biological Chemistry 2016, 291, 6696–6705.

(18) Li, P.; Banjade, S.; Cheng, H.-C.; Kim, S.; Chen, B.; Guo, L.; Llaguno, M.; Hollingsworth, J. V.; King, D. S.; Banani, S. F., et al. Nature 2012, 483, 336–340.

(19) Dunker, A. K.; Lawson, J. D.; Brown, C. J.; Williams, R. M.; Romero, P.; Oh, J. S.; Oldfield, C. J.; Campen, A. M.; Ratliff, C. M.; Hipps, K. W., et al. Journal of molecular graphics and modelling 2001, 19, 26–59.

(20) Uversky, V. N.; Oldfield, C. J.; Dunker, A. K. Journal of Molecular Recognition: An Interdisciplinary Journal 2005, 18, 343–384.

(21) Van Der Lee, R.; Buljan, M.; Lang, B.; Weatheritt, R. J.; Daughdrill, G. W.; Dunker, A. K.; Fuxreiter, M.; Gough, J.; Gsponer, J.; Jones, D. T., et al. Chemical reviews 2014, 114, 6589.

(22) Nott, T. J.; Petsalaki, E.; Farber, P.; Jervis, D.; Fussner, E.; Plochowietz, A.; Craggs, T. D.; Bazett-Jones, D. P.; Pawson, T.; Forman-Kay, J. D., et al. Molecular cell 2015, 57, 936–947.

(23) Brangwynne, C. P.; Tompa, P.; Pappu, R. V. Nature Physics 2015, 11, 899–904.

(24) Burke, K. A.; Janke, A. M.; Rhine, C. L.; Fawzi, N. L. Molecular cell 2015, 60, 231–241.

(25) Elbaum-Garfinkle, S.; Kim, Y.; Szczepaniak, K.; Chen, C. C.-H.; Eckmann, C. R.; Myong, S.; Brangwynne, C. P. Proceedings of the National Academy of Sciences 2015, 112, 7189–7194.

(26) Rekhi, S.; Garcia, C. G.; Barai, M.; Rizuan, A.; Schuster, B. S.; Kiick, K. L.; Mittal, J. Nature chemistry 2024, 16, 1113–1124.

(27) Lin, Y.-H.; Forman-Kay, J. D.; Chan, H. S. Physical review letters 2016, 117, 178101.

(28) Martin, E. W.; Holehouse, A. S.; Peran, I.; Farag, M.; Incicco, J. J.; Bremer, A.; Grace, C. R.; Soranno, A.; Pappu, R. V.; Mittag, T. Science 2020, 367, 694–699.

(29) Schuster, B. S.; Dignon, G. L.; Tang, W. S.; Kelley, F. M.; Ranganath, A. K.; Jahnke, C. N.; Simpkins, A. G.; Regy, R. M.; Hammer, D. A.; Good, M. C., et al. Proceedings of the National Academy of Sciences 2020, 117, 11421–11431.

(30) Singh, A.; Ukperaj, A. I.; Porto, G. F.; Dignon, G. L. PLOS Computational Biology 2026, 22, e1014462.

(31) Cai, H.; Vernon, R. M.; Forman-Kay, J. D. Biomolecules 2022, 12, 1131.

(32) Cohan, M. C.; Shinn, M. K.; Lalmansingh, J. M.; Pappu, R. V. Journal of molecular biology 2022, 434, 167373.

(33) Kilgore, H. R.; Chinn, I.; Mikhael, P. G.; Mitnikov, I.; Van Dongen, C.; Zylberberg, G.; Afeyan, L.; Banani, S. F.; Wilson-Hawken, S.; Lee, T. I., et al. Science 2025, 387, 1095–1101.

(34) Pak, C. W.; Kosno, M.; Holehouse, A. S.; Padrick, S. B.; Mittal, A.; Ali, R.; Yunus, A. A.; Liu, D. R.; Pappu, R. V.; Rosen, M. K. Molecular cell 2016, 63, 72–85.

(35) Ruff, K. M.; King, M. R.; Ying, A. W.; Liu, V.; Pant, A.; Lieberman, W. E.; Shinn, M. K.; Su, X.; Kadoch, C.; Pappu, R. V. Cell 2026, 189, 323–342.

(36) Das, R. K.; Pappu, R. V. Proceedings of the National Academy of Sciences 2013, 110, 13392–13397.

(37) Sawle, L.; Ghosh, K. The Journal of chemical physics 2015, 143.

(38) Zheng, W.; Dignon, G.; Brown, M.; Kim, Y. C.; Mittal, J. The journal of physical chemistry letters 2020, 11, 3408.

(39) Lin, Y.-H.; Chan, H. S. Biophysical journal 2017, 112, 2043–2046.

(40) Statt, A.; Casademunt, H.; Brangwynne, C. P.; Panagiotopoulos, A. Z. The Journal of chemical physics 2020, 152.

(41) Weiner, B. G.; Pyo, A. G.; Meir, Y.; Wingreen, N. S. PLoS computational biology 2021, 17, e1009748.

(42) Alshareedah, I.; Borcherds, W. M.; Cohen, S. R.; Singh, A.; Posey, A. E.; Farag, M.; Bremer, A.; Strout, G. W.; Tomares, D. T.; Pappu, R. V., et al. Nature Physics 2024, 20, 1482–1491.

(43) Banani, S. F.; Rice, A. M.; Peeples, W. B.; Lin, Y.; Jain, S.; Parker, R.; Rosen, M. K. Cell 2016, 166, 651–663.

(44) Kuznetsova, K.; Scheremetjew, M.; Yin, J.; Moon, H.; Vargas, D. A.; Hadarovich, A.; Lewis, N.; Hoege, C.; Chow, C. F. W.; Kuster, D., et al. Nucleic Acids Research 2026, 54, D375–D382.

(45) Saha, S.; Weber, C. A.; Nousch, M.; Adame-Arana, O.; Hoege, C.; Hein, M. Y.; Osborne-Nishimura, E.; Mahamid, J.; Jahnel, M.; Jawerth, L., et al. Cell 2016, 166, 1572–1584.

(46) Youn, J.-Y.; Dyakov, B. J.; Zhang, J.; Knight, J. D.; Vernon, R. M.; Forman-Kay, J. D.; Gingras, A.-C. Molecular cell 2019, 76, 286–294.

(47) Guzikowski, A. R.; Chen, Y. S.; Zid, B. M. Wiley Interdisciplinary Reviews: RNA 2019, 10, e1524.

(48) Xing, W.; Muhlrad, D.; Parker, R.; Rosen, M. K. Elife 2020, 9, e56525.

(49) Welles, R. M.; Sojitra, K. A.; Garabedian, M. V.; Xia, B.; Wang, W.; Guan, M.; Regy, R. M.; Gallagher, E. R.; Hammer, D. A.; Mittal, J., et al. Nature chemistry 2024, 16, 1062–1072.

(50) Lin, Y.-H.; Brady, J. P.; Forman-Kay, J. D.; Chan, H. S. New Journal of Physics 2017, 19, 115003.

(51) Dignon, G. L.; Best, R. B.; Mittal, J. Annual review of physical chemistry 2020, 71, 53–75.

(52) Chew, P. Y.; Joseph, J. A.; Collepardo-Guevara, R.; Reinhardt, A. Chemical Science 2023, 14, 1820– 1836.

(53) Ginell, G. M.; Emenecker, R. J.; Lotthammer, J. M.; Keeley, A. T.; Plassmeyer, S. P.; Razo, N.; Usher, E. T.; Pelham, J. F.; Holehouse, A. S. Science 2025, 388, eadq8381.

(54) Hunter, K.; Brandt, T.; Guadalupe, K.; Kolamunna, K. C.; Lotthammer, J. M.; Shamoon, N. M.; Niblo, J. K.; Nicholson, B.; Day, L. M.; Martinez, A., et al. Nature 2026, 1–10.

(55) Wessén, J.; De La Cruz, N.; Lyons, H.; Chan, H. S.; Sabari, B. R. Communications Chemistry 2025, 8, 304.

(56) Villegas, J. A.; Levy, E. D. Protein Science 2022, 31, _eprint: https://onlinelibrary.wiley.com/doi/pdf/10.1002/pro.436e4361.

(57) Morton, W.; Vácha, R. bioRxiv 2025, 2025–12.

(58) Regy, R. M.; Thompson, J.; Kim, Y. C.; Mittal, J. Protein Science 2021, 30, 1371–1379.

(59) Dignon, G. L.; Zheng, W.; Kim, Y. C.; Best, R. B.; Mittal, J. PLoS computational biology 2018, 14, e1005941.

(60) Farag, M.; Cohen, S. R.; Borcherds, W. M.; Bremer, A.; Mittag, T.; Pappu, R. V. Nature communications 2022, 13, 7722.

(61) Dignon, G. L.; Zheng, W.; Best, R. B.; Kim, Y. C.; Mittal, J. Proceedings of the National Academy of Sciences 2018, 115, 9929–9934.

(62) Urry, D. W.; Gowda, D. C.; Parker, T. M.; Luan, C.-H.; Reid, M. C.; Harris, C. M.; Pattanaik, A.; Harris, R. D. Biopolymers: Original Research on Biomolecules 1992, 32, 1243–1250.

(63) Eastman, P.; Swails, J.; Chodera, J. D.; McGibbon, R. T.; Zhao, Y.; Beauchamp, K. A.; Wang, L.-P.; Simmonett, A. C.; Harrigan, M. P.; Stern, C. D., et al. PLoS computational biology 2017, 13, e1005659.

(64) Ashbaugh, H. S.; Hatch, H. W. Journal of the American Chemical Society 2008, 130, 9536–9542.

(65) Debye, P.; Hückel, E. Phys. Z 1923, 24, 185–206.

(66) Silmore, K. S.; Howard, M. P.; Panagiotopoulos, A. Z. Molecular Physics 2017, 115, 320–327.

